# IRTKS promotes Tir membrane insertion for intimate bacterial attachment and subsequent pedestal formation

**DOI:** 10.64898/2026.05.21.727007

**Authors:** Julissa Burgos-Rivera, Faiz Z. Maredia, Carina I. Roman-Aquino, Matthew J. Tyska

## Abstract

Enterohemorrhagic *Escherichia coli* (EHEC) is a foodborne pathogen that causes bloody diarrhea and hemolytic uremic syndrome by disrupting the intestinal brush border. During infection, EHEC injects the transmembrane virulence factor Tir into enterocytes; upon insertion into the apical membrane, this factor mediates bacterial attachment and drives formation of actin-rich pedestals needed for colonization. How Tir is inserted into the host plasma membrane remains unclear. Here, we investigated the role of brush border resident IRTKS, a Tir- and membrane-binding protein, in this process. Using multiple IRTKS gain- and loss-of-function models, we analyzed pedestal organization and component localization. Whereas canonical models position IRTKS downstream of Tir as a scaffolding link to F-actin, we found that perturbing IRTKS disrupted the distribution and abundance of Tir. Moreover, ectopic IRTKS expression enhanced Tir membrane insertion in the absence of other virulence factors. We conclude that IRTKS functions early in pedestal formation to promote Tir accumulation in the plasma membrane and in turn, facilitate bacterial attachment.

**SUMMARY:** EHEC attaches to intestinal epithelial cells using injected virulence factor Tir, which forms actin pedestals and promotes bacterial colonization. We found that host protein IRTKS promotes Tir accumulation in the plasma membrane, to facilitate intimate bacterial attachment and pedestal formation.

## INTRODUCTION

The functional surface of the small intestine consists of a diverse collection of epithelial cell types, which form a monolayer that supports selective uptake of nutrients, while simultaneously serving as a barrier to luminal contents and the microbes that reside in this space. To maximize nutrient uptake, enterocytes, the primary solute transporting cell type in the small intestine, build an array of microvilli on their apical surface, which increases absorptive surface area up to ∼15-fold (Crawley et al., 2014; Helander and Fandriks, 2014; Louvard et al., 1992). Each microvillus is supported by ∼20-40 bundled, parallel actin filaments, which extend their barbed ends toward the distal tip and pointed ends down into a sub-apical terminal web (Mooseker and Tilney, 1975; Ohta et al., 2012). During enterocyte differentiation, the core actin bundles in nascent microvilli exhibit treadmilling, powered by F-actin turnover, which drives their motion across the apical surface, collisions between neighboring protrusions, and adhesive consolidation into large, organized clusters (Loomis et al., 2003; Meenderink et al., 2019; Tyska and Mooseker, 2002). As clusters increase in size, they eventually fill the enterocyte apical surface forming the ‘brush border’. Recently, it was shown that microvilli at the periphery of the brush border form adhesive contacts with microvilli on neighboring cells (Cencer et al., 2024). Together with canonical junctional complexes that are found more basally (e.g. tight and adherence junctions), these interactions may contribute to the structural barrier function that limits luminal pathogens and toxins from entering peripheral tissues (Suzuki, 2013).

Despite the structural barrier presented by the intestinal epithelium, several enteric pathogens have evolved unique infectious mechanisms to survive and colonize the small intestine, without gaining direct access to peripheral tissues (Goosney et al., 1999; Rodrigues et al., 2025). A well-studied example is Enterohemorrhagic *Escherichia coli* (EHEC) O157:H7, which emerged as a foodborne pathogen in 1982, when it was first isolated from two outbreaks linked to contaminated hamburger meat in the United States (Riley et al., 1983). EHEC continues to cause multiple disease outbreaks worldwide each year, resulting in bloody diarrhea and hemolytic uremic syndrome, a common cause of pediatric renal failure (Griffin and Tauxe, 1991; Majowicz et al., 2014). These symptoms are caused by EHEC’s production of Shiga toxin, an AB_5_ toxin, that targets the colon and kidney, and from the focal disruption of the intestinal epithelium through the formation of attaching and effacing (A/E) lesions on the apical surface of enterocytes (Joseph et al., 2020; Obrig, 2010; Paton and Paton, 1998; Robinson et al., 2006; Scotland et al., 1985). A/E lesions are defined by the effacement of enterocyte microvilli, intimate attachment of extracellular EHEC to the effaced apical surface, and reorganization of the host cytoskeletal machinery to form dynamic actin-rich structures known as pedestals (Frankel et al., 1998). Previous work on EHEC and Enteropathogenic *E. coli* (EPEC), another human A/E pathogen, revealed the importance of intimate attachment and pedestal formation for colonization and disease outcome in animal and cell culture models (Donnenberg et al., 1993b; Knutton et al., 1989; Marches et al., 2000; McKee et al., 1995; Ritchie et al., 2008; Ritchie et al., 2003; Stavric et al., 1993; Tzipori et al., 1995), and human volunteers (Donnenberg et al., 1993a; Tacket et al., 2000). Despite progress toward understanding the function of A/E lesions and pedestals, mechanisms that control EHEC intimate attachment for pedestal formation remain unclear.

A/E pathogens are distinguished by the presence of a pathogenicity island within their genome known as locus of enterocyte effacement (LEE), which encodes the components that drive intimate bacterial attachment and activate host actin remodeling for pedestal formation (Elliott et al., 1998; Perna et al., 1998). During initial attachment, EHEC adheres to the host enterocyte using its flagella and pilus/fimbrial adhesins, although the precise mechanisms remain undefined (Croxen et al., 2013; Mahajan et al., 2009; McWilliams and Torres, 2014; Rendon et al., 2007; Torres et al., 2002; Xicohtencatl-Cortes et al., 2007). EHEC then uses its Type III Secretion System (T3SS) to secrete bacterial effectors into the host cell, which in turn drive the formation of A/E lesions (Jarvis and Kaper, 1996). A key secreted effector is Translocated intimin receptor (Tir), previously known as Hp90, which is required for intimate bacterial attachment and for initiating host cell actin assembly machinery for pedestal formation (Kenny et al., 1997; Rosenshine et al., 1992). In Tir-mediated attachment, Tir is inserted in the apical membrane as a hairpin-loop flanked by two transmembrane domains through an undefined mechanism (Campellone et al., 2006; DeVinney et al., 1999). The central extracellular domain of Tir then binds to intimin, an LEE-encoded bacterial outer membrane adhesin (DeVinney et al., 1999; Liu et al., 1999). The Tir-intimin interaction supports the intimate attachment of EHEC to the host cell, which is required for colonization (Battle et al., 2014; Ritchie et al., 2003; Tzipori et al., 1995; Yu and Kaper, 1992). To initiate pedestal formation, the intracellular C-terminus of Tir recruits and binds to the Inverse-Bin-Amphiphysin-Rvs (I-BAR) domain of host proteins Insulin Receptor Tyrosine Kinase Substrate (IRTKS), the primary adaptor in this pathway, as well as the related protein Insulin Receptor Tyrosine Kinase Substrate p53 (IRSp53) (Allen-Vercoe et al., 2006; Campellone et al., 2006; de Groot et al., 2011; Vingadassalom et al., 2009; Weiss et al., 2009; Yi and Goldberg, 2009). IRTKS serves as a molecular bridge to secreted effector *E. coli* secreted protein F in prophage U (EspF_U_) (also known as TccP), which is not LEE-encoded, via its SRC Homology 3 (SH3) domain (Aitio et al., 2010; Campellone et al., 2004; Garmendia et al., 2004; Ritchie et al., 2008). The resulting Tir-IRTKS-EspF_U_ complex that forms beneath adherent bacteria triggers actin pedestal assembly. EspF_U_ promotes actin polymerization by recruiting N- WASP, a host actin nucleation promoting factor, which then activates the Arp2/3 complex to nucleate branched actin filaments that form the cytoskeletal core of the pedestal (Campellone et al., 2008; Cheng et al., 2008; Sallee et al., 2008) (Figure 1A). Beyond mediating strong attachment to the cell surface, pedestal motility aids in cell-to-cell spread to enhance EHEC colonization throughout the small intestine (Shaner et al., 2005; Velle and Campellone, 2017).

**Figure 1:**
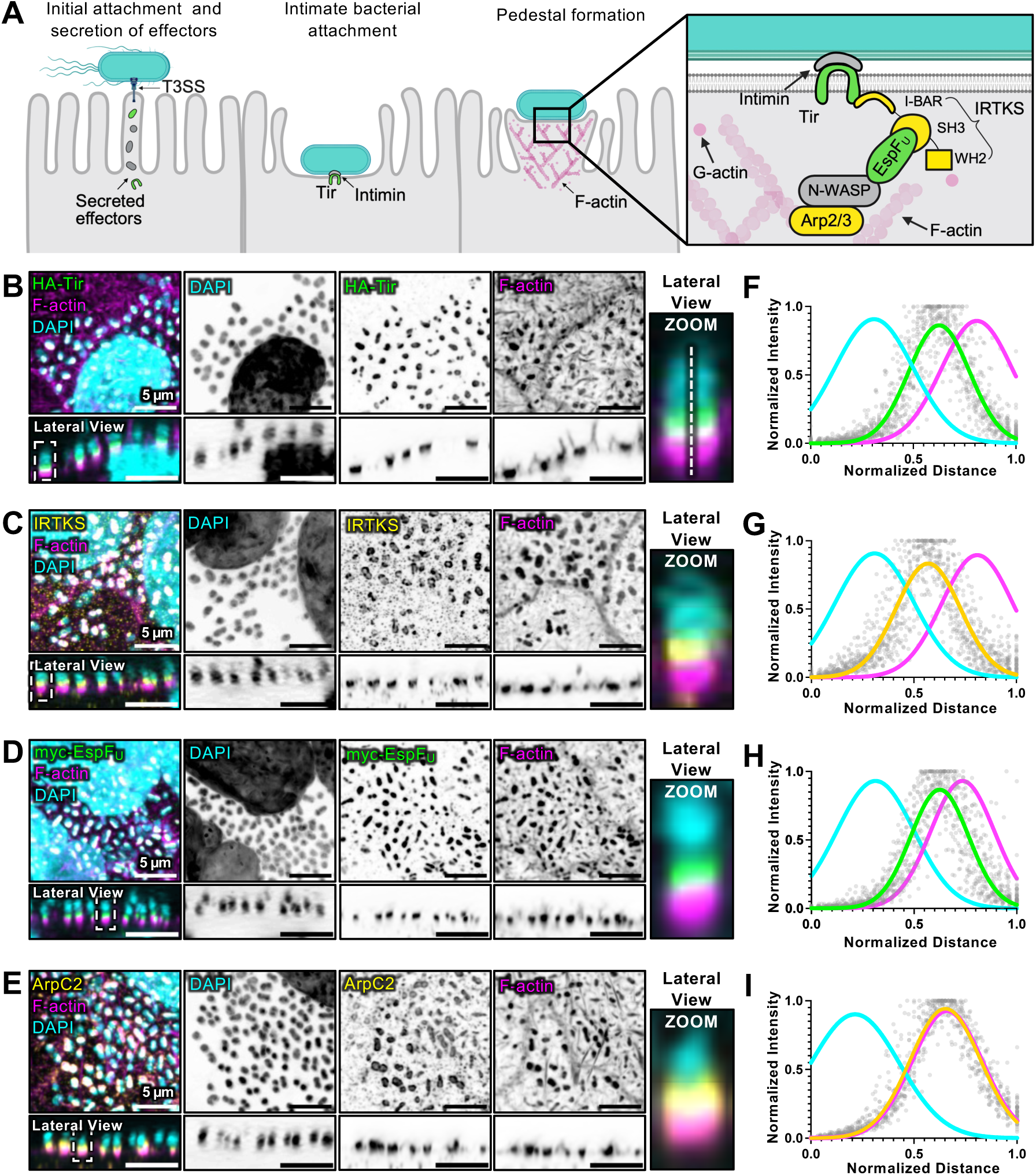
IRTKS and its interactors are enriched in pedestals formed on the apical surface of CACO-2BBE cells. (A) Current model of EHEC Tir-mediated attachment and pedestal formation. Tir (green) is translocated to the plasma membrane, where it binds intimin for intimate bacterial attachment and initiates the recruitment of actin-associated proteins for pedestal formation. IRTKS (yellow) binds to Tir and EspF_U_ (green) via its I-BAR and SH3 domain, respectively. EspF_U_ recruits nucleating-promoting factor, N-WASP, to activate the Arp2/3 complex (yellow) for actin pedestal assembly. (B-E) Maximum intensity projections of confocal volumes and lateral views (below) showing the localization of (B) HA-Tir (green), (C) IRTKS (yellow), (D) myc-EspF_U_ (green), and (E) ArpC2 (yellow) and co-stained with DAPI (cyan) and F-actin (magenta) in KC12+EspF_U_ infected CACO-2BBE. Zoom inset in B shows a representative line scan used for quantification. (F-I) Line scan plots of individual pedestal-forming bacteria show the intensity distribution and localization of proteins-of-interest in relation to DAPI and F-actin; 0, distal end of bacteria; 1, basal end of F-actin signal. Grey points on the plot represent intensity signals throughout the pedestal structure for the protein-of-interest. Solid lines represent Gaussian fits to the DAPI, protein-of-interest, and F-actin signals in each case; n = 45 pedestals from three experimental replicates. Scale bars represent 5 µm.

Despite the well-established role of Tir in bacterial attachment and pedestal formation, molecular mechanisms that drive the insertion and accumulation of Tir in the apical plasma membrane remain poorly understood. Based on its hairpin-loop topology, previous studies suggested that insertion takes place via a mechanism distinct from the canonical ER-dependent biosynthetic pathway used by eukaryotic transmembrane proteins (Kenny et al., 1997). Supporting this, an *in vitro* study showed that purified Tir can associate with and insert into lipid vesicles (Race et al., 2006). However, other work showed that disruption of the Golgi apparatus with brefeldin A blocks Tir localization to the plasma membrane, suggesting that host secretory trafficking might also contribute to Tir accumulation (Mao et al., 2017). Another possible mechanism involves host factors, for example IRTKS, which contains an N-terminal I-BAR domain that binds to Tir and normally associates with the apical plasma membrane. IRTKS was first identified in studies of insulin signaling and then as member of the I-BAR protein family involved in driving outward membrane curvature for protrusion formation (Millard et al., 2007; Zhao et al., 2011). More recently, IRTKS was shown to localize to the tips of growing microvilli, and its I-BAR domain was found to contribute to tip targeting (Gaeta et al., 2021; Postema et al., 2018). In the context of EHEC infection, previous studies suggested that IRTKS functions as a scaffold, downstream of Tir, linking to EspF_U_ for pedestal formation (Vingadassalom et al., 2009). Whether IRTKS also plays roles in recruiting Tir to or insertion into the apical plasma membrane, upstream of pedestal formation, remains unclear.

In this study, we sought to test the hypothesis that, through its Tir- and membrane-binding I-BAR domain, IRTKS promotes Tir accumulation at the plasma membrane, upstream of pedestal formation. In our experiments, we leveraged several new IRTKS gain- and loss-of-function models, coupled with high resolution microscopy to assay the localization and organization of pedestal components in response to these perturbations. Although established models place IRTKS downstream of Tir as a scaffolding link to EspF_U_ and the Arp2/3 actin polymerization machinery, we unexpectedly found that perturbing IRTKS disrupted both the distribution and abundance of Tir. We also observed that ectopic IRTKS expression directly enhanced Tir membrane insertion in the absence of other virulence factors. Based on our findings, we conclude that, in addition to its EspF_U_ scaffolding role, IRTKS functions early in EHEC’s infection pathway to promote Tir accumulation in the plasma membrane and in turn, facilitate intimate bacterial attachment.

## RESULTS

### IRTKS and its interactors exhibit a conserved, stratified distribution in pedestals

As a first step toward clarifying the mechanism of Tir insertion into the apical plasma membrane, we sought to define the localization of its binding partner, IRTKS, along with other key bacterial and host cell factors that are recruited to pedestals (Fig. 1A). As a target cell culture model for these experiments, we chose the CACO-2BBE cell line, human colonic epithelial cells that form a well-organized monolayer and acquire enterocyte-like biochemical and morphological features after several weeks in culture (Peterson et al., 1993; Peterson and Mooseker, 1992; Peterson and Mooseker, 1993). CACO-2BBE cells assemble an apical brush border that closely recapitulates the *in vivo* target encountered by A/E pathogens. As a model pathogenic microbe, we used KC12+EspF_U_, an EPEC strain engineered to express EHEC intimin, HA-tagged Tir (HA-Tir), and myc-tagged EspF_U_ (myc-EspF_U_), the complement of EHEC proteins that are sufficient for pedestal formation. As a previously established model for EHEC infection, this strain allows the visualization of both bacterial effectors, Tir and EspF_U_ (Campellone et al., 2004; Heath et al., 2011; Velle and Campellone, 2017; Velle and Campellone, 2018; Vingadassalom et al., 2010). CACO-2BBE cells were grown to 21 days post-confluency (DPC) to form highly differentiated monolayers, infected with KC12+EspF_U_, fixed and stained for pedestal components, and then imaged with confocal microscopy. Pedestals exhibited robust recruitment of HA-Tir, IRTKS, myc-EspF_U_, ArpC2, a subunit of the Arp2/3 complex, and F-actin to bacterial attachment sites (Fig. 1B–1E). To quantify the distribution of these factors in individual pedestals, we performed line scan analysis on projected lateral views of the apical surface (Fig. 1F–1I). Line scans analysis of pedestals revealed that HA-Tir, IRTKS, and myc-EspF_U_ exhibit strong overlap with each other, accumulating in a layered manner between the bacterial and F-actin signals (Fig. 1F–1H). In contrast, ArpC2 demonstrated robust colocalization with F-actin at the basal end of the pedestal (Fig. 1I).

To determine if the localization pattern of pedestal components observed here is organized by the infecting pathogen or, instead, by the nature of the host cell target, we also used KC12+EspF_U_ to infect HeLa cells, a non-polarizing human culture model that has been extensively employed in previous studies of pedestal formation (Allen-Vercoe et al., 2006; Battle et al., 2014; Campellone et al., 2006; Campellone et al., 2008; Campellone et al., 2004; DeVinney et al., 1999; Garmendia et al., 2004; Vingadassalom et al., 2009). Analysis of HA-Tir, IRTKS, myc-EspF_U_, ArpC2, and F-actin localization in pedestals on the surface of HeLa cells revealed a distribution and stratified organization similar to that observed on CACO-2BBE cells (Fig. 2). Together, these findings reveal that, independent of the target host cell context, pedestals assembled by KC12+EspF_U_ exhibit a robust, layered organization of components.

**Figure 2:**
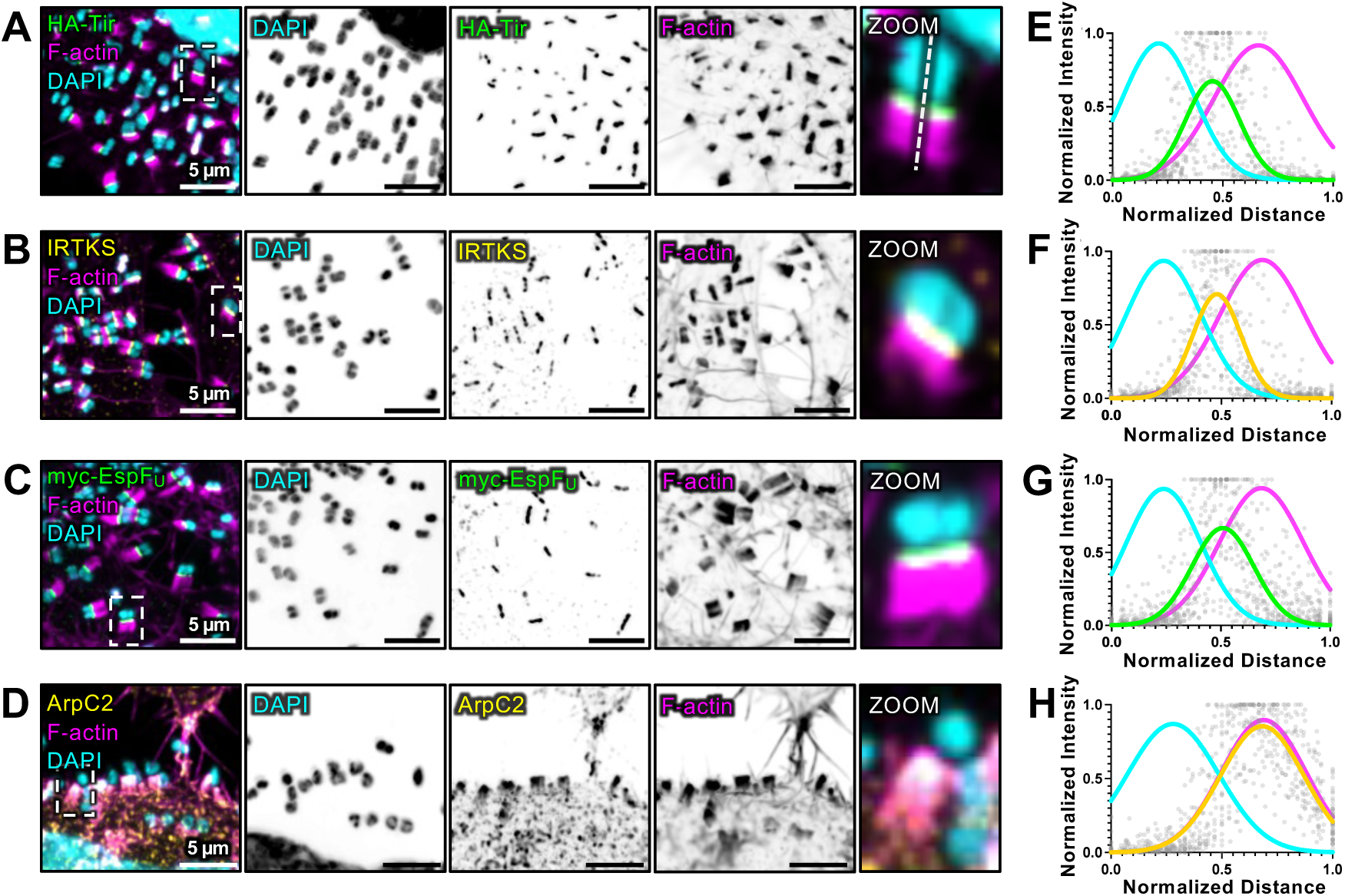
Key proteins exhibit a conserved, stratified distribution in pedestals on the surface of HeLa cells. (A-D) Maximum intensity projections of confocal image volumes showing the localization of (A) HA-Tir (green), (B) IRTKS (yellow), (C) myc-EspF_U_ (green), and (D) ArpC2 (yellow) in pedestals co-stained with DAPI (cyan) and F-actin (magenta) in KC12+EspF_U_ infected HeLa cells. Zoom inset in A shows a representative line scan used for quantification. (E-H) Line scan plots of individual pedestal-forming bacteria showing the intensity distribution and localization of each protein-of-interest in relation to DAPI and F-actin; 0, distal end of bacteria; 1, basal end of F-actin signal. Grey points on the plot represent intensity signals throughout the pedestal structure for the protein-of-interest. Solid lines represent Gaussian fits to the DAPI, protein-of-interest, and F-actin signals in each case; n = 45 pedestals were quantified from three experimental replicates. Scale bars represent 5 µm.

### Overexpression of IRTKS disperses Tir and increases bacterial attachment in an I-BAR dependent manner

The strong colocalization of HA-Tir and IRTKS observed in our pedestal imaging experiments is expected given that these proteins are interacting partners (Vingadassalom et al., 2009). Indeed, previous biochemical studies established that the C-terminal cytoplasmic tail of Tir binds directly to the N-terminal I-BAR domain of IRTKS (de Groot et al., 2011). Because the I-BAR also binds directly to membranes, IRTKS is a promising candidate for promoting Tir recruitment to the plasma membrane (Millard et al., 2007; Postema et al., 2018; Zhao et al., 2011). From this perspective, we sought to investigate the role of IRTKS and its I-BAR domain in Tir localization and Tir-mediated bacterial attachment. Here, we used KC12+EspF_U_ to infect HeLa cells that were first transfected with one of the following EGFP-tagged constructs: full-length IRTKS, an I-BAR deletion mutant (ΔI-BAR), the I-BAR domain only (I-BAR), or a variant of full-length IRTKS that blocks ligand binding in the SH3 domain (SH3*) but retains an intact I-BAR domain (Fig. 3A). Expression of each construct was confirmed by Western blotting using anti-IRTKS and anti-EGFP probes (Fig. 3B–3C). Whereas control cells exhibited a tightly focused Tir signal beneath individual bacterial attachment sites, IRTKS overexpression led to a striking dispersion of Tir across the cell surface (Fig. 3D–3E), as indicated by a significant increase in the percentage of the surface covered by HA-Tir (Fig. 3I). To understand how Tir dispersion impacted bacterial attachment, we quantified the Lipopolysaccharide (LPS) signal volume as a metric for the total amount of bound bacteria. This measurement revealed that IRTKS overexpression significantly increased bacterial attachment (Fig. 3D-3E, 3J). We also observed a strong trending increase in HA-Tir sum intensity, suggesting that total levels of Tir were elevated (Fig. 3K). Analysis of truncated and mutated IRTKS variants further revealed that the presence of the I-BAR domain was required for both Tir dispersion and the associated increase in bacterial attachment (Fig. 3F-3H, 3I–3J). Cells overexpressing the ΔI-BAR mutant showed HA-Tir dispersion and LPS signal levels similar to untransfected controls (Fig. 3D, 3F, 3I–3J), whereas constructs that contained an intact I-BAR domain (I-BAR and SH3*) trended toward higher levels of Tir dispersion and increased bacterial adherence (Fig. 3G-H, 3I–3J). Together, these data indicate that overexpression of IRTKS or constructs containing the I-BAR domain, leads to Tir dispersion and an increase in bacterial adherence; they also suggest that IRTKS may promote Tir enrichment at the plasma membrane for bacterial adherence.

**Figure 3:**
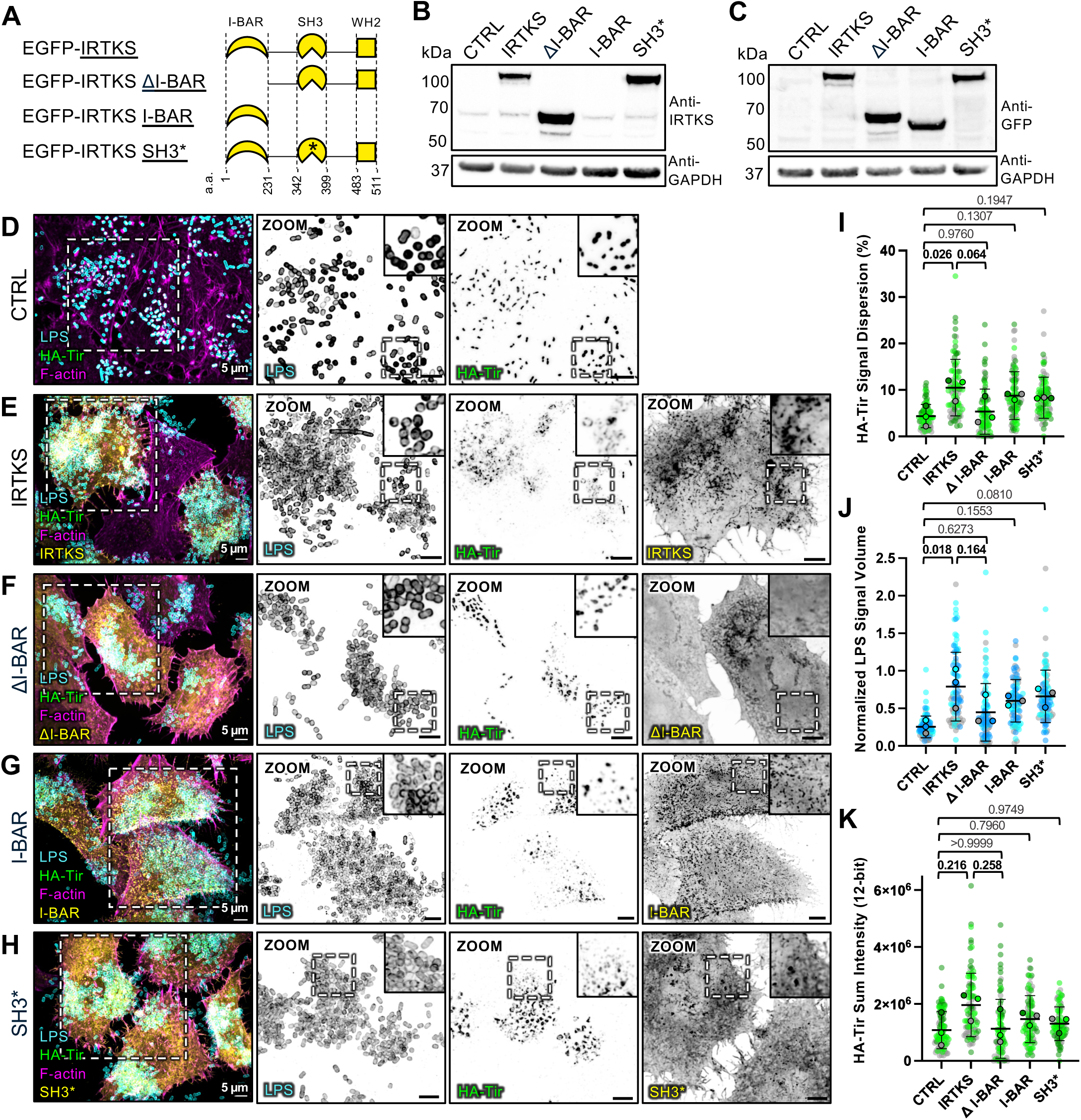
Overexpression of IRTKS disperses Tir and increases bacterial attachment in an I-BAR dependent manner. (A) Domain diagrams of IRTKS overexpression constructs used in this study; * refers to W378K/W391K point mutations within the SH3 domain; a.a. refers to the amino acid numbers. Underlines refer to construct names used throughout this study. (B-C) Western blot analysis of EGFP-IRTKS construct expression in HeLa cells. To visualize all constructs, both (B) IRTKS and (C) GFP antibodies were used, as the IRTKS antibody targets a peptide sequence within the SH3 domain. (D-H) Maximum intensity projections of confocal image volumes of KC12+EspF_U_ infected HeLa cells (D) Control (untransfected) and overexpressing (E) EGFP-IRTKS, (F) EGFP-IRTKS ΔI-BAR, (G) EGFP-IRTKS I-BAR, or (H) EGFP-IRTKS SH3* (yellow) and stained for LPS (cyan), HA-Tir (green), and F-actin (magenta). (I) Quantification of percentage HA-Tir signal dispersion, calculated as the thresholded HA-Tir fluorescence signal area normalized to total cell area; 30 cells per condition were quantified per experimental replicate, with n = three experimental replicates. (J) Quantification of thresholded LPS fluorescence signal volume per cell, normalized to total cell area; 30 cells per condition were quantified per experimental replicate, with n = three experimental replicates. (K) Sum of HA-Tir 12-bit intensity values per cell from thresholded fluorescence signal; 30 cells per condition were quantified per experimental replicate, with n = three experimental replicates. (I-K) Data are shown as superplots, where transparent circles represent individual cell measurements color-coded by experimental replicate, and opaque outlined circles show the means of each experimental replicate. Statistical comparisons were performed on experimental replicate means. All p-values were calculated with an ordinary one-way ANOVA with multiple comparisons; exact p-values are shown above brackets. Error bars represent the mean ± SD of the technical replicates. Scale bars represent 5 µm.

### Overexpression of IRTKS mislocalizes EspF_U_ and disrupts pedestal formation

Tir insertion into the plasma membrane and intimate bacterial attachment, which are both impacted by IRTKS overexpression, are upstream of EspF_U_ recruitment and actin pedestal assembly (Campellone et al., 2004; Vingadassalom et al., 2009). Thus, we next assessed EspF_U_ localization and pedestal formation in KC12+EspF_U_ infected cells ectopically expressing IRTKS variants. Whereas untransfected control cells demonstrated strong, focused enrichment of myc-EspF_U_ in F-actin dense pedestals (Fig. 4A), cells overexpressing IRTKS exhibited severely mislocalized myc-EspF_U_ (Fig. 4A-4B, 4K) with myc-EspF_U_ dispersed to regions distinct from Tir (Fig. S1). Lateral views of single cells also indicated that dispersed myc-EspF_U_ accumulated near the plasma membrane (Fig. 4F, 4G, arrows). We also detected a significant increase in myc-EspF_U_ total sum intensity per cell with IRTKS overexpression (Fig. 4L), which likely reflects the increased number of adherent bacteria under these conditions (Fig. 3J). Despite the increase in myc-EspF_U_, we observed a striking reduction in pedestal formation efficiency (Fig. 4M).

**Figure 4:**
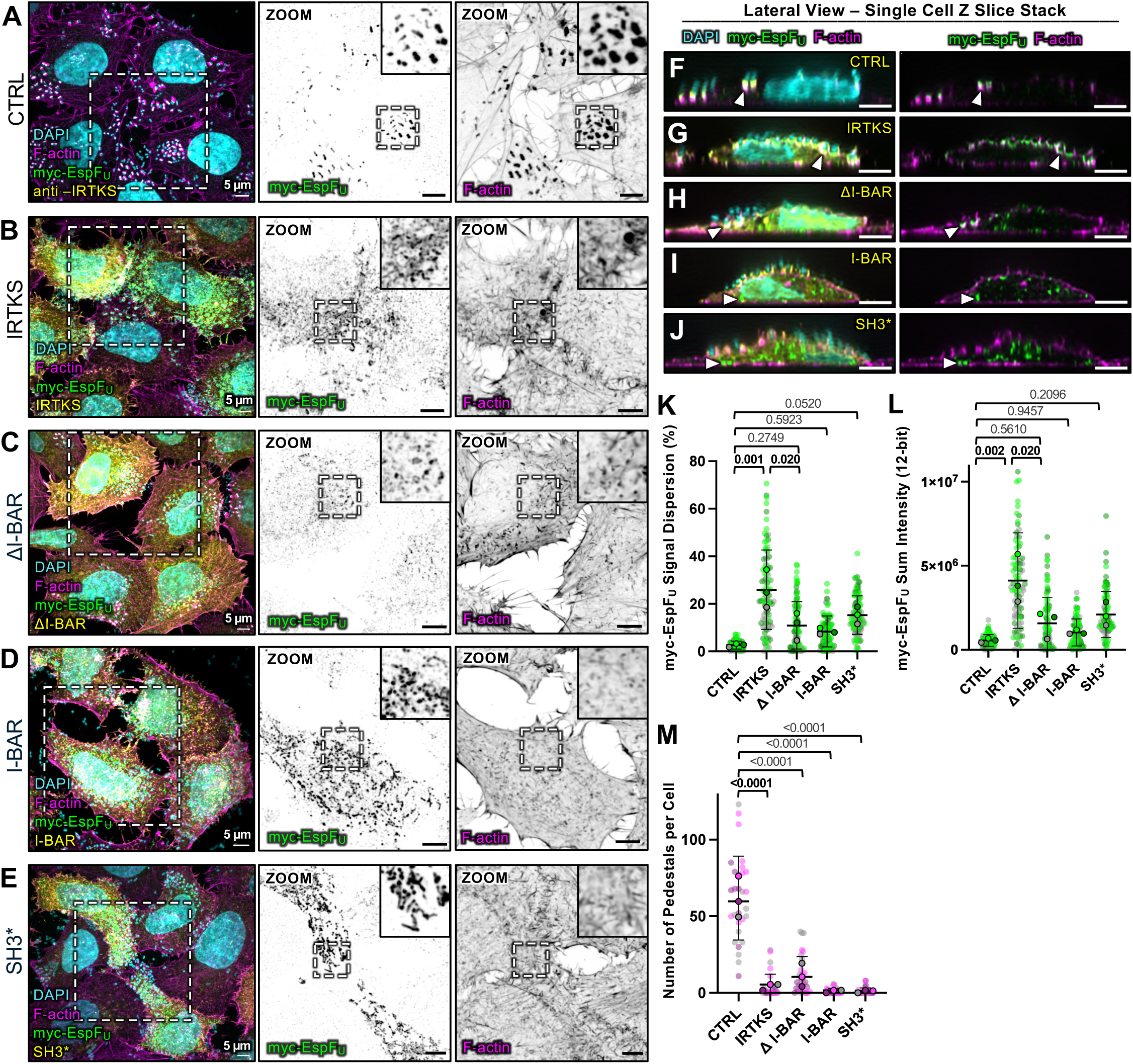
Overexpression of IRTKS mislocalizes EspF_U_ and disrupts pedestal formation. (A-E) Maximum intensity projections of confocal image volumes of KC12+EspF_U_ infected HeLa cells. (A) Control (untransfected) cells stained for endogenous IRTKS, and cells overexpressing (B) EGFP–IRTKS, (C) EGFP–IRTKS ΔI-BAR, (D) EGFP–IRTKS I-BAR, or (E) EGFP–IRTKS SH3* (yellow). All samples were stained for DAPI (cyan), myc-EspF_U_ (green), and F-actin (magenta). (F-J) Lateral view of single cell z-slices showing endogenous (F) IRTKS, and overexpressed constructs (G) EGFP-IRTKS, (H) EGFP-IRTKS ΔI-BAR, (I) EGFP-IRTKS I-BAR, or (J) EGFP-IRTKS SH3*. Arrows indicate the localization of myc-EspF_U_ in the cell. (K) Quantification of the percentage of myc-EspF_U_ signal dispersion, calculated as thresholded myc-EspF_U_ fluorescence signal area normalized to the total cell area; 30 cells per condition were quantified per experimental replicate, with n = three experimental replicates. (L) Sum of 12-bit myc-EspF_U_ intensity values per cell; 30 cells per condition were quantified per experimental replicate, with n = three experimental replicates. (M) Quantification of the number of pedestals per cell. Pedestal formation efficiency was determined by the intense F-actin signal beneath the bacteria; 10 cells per condition were quantified per experimental replicate, with n = three experimental replicates. (K-M) Data are shown as superplots, where transparent circles represent individual cell measurements color-coded by experimental replicate, and opaque outlined circles show the means of each experimental replicate. Statistical comparisons were performed on experimental replicate means. All p-values were calculated with an ordinary one-way ANOVA with multiple comparisons; exact p-values are shown above brackets. Error bars represent the mean ± SD of the technical replicates. Scale bars represent 5 µm.

For each of the three mutant IRTKS constructs (Fig. 3A), myc-EspF_U_ signal dispersion and sum intensity trended upward, although levels were not significantly different from controls (Figure 4C-4E,4K–4L). Surprisingly, in constructs lacking a functional SH3 domain (I-BAR and SH3*), myc-EspF_U_ mislocalized intracellularly (Fig. 4I–4J, arrows). Pearson’s correlation analysis showed strong colocalization with TOMM20, a mitochondrial marker (Fig. S2), suggesting that myc-EspF_U_ is mislocalized when the SH3 domain is nonfunctional. Furthermore, like full length IRTKS, these variants uniformly abolished pedestal formation (Fig. 4M). These experiments reveal that although IRTKS overexpression disperses Tir and leads to higher levels of bacterial attachment (Fig. 3), these perturbations do not lead to more extensive EspF_U_ recruitment for pedestal formation. Instead, IRTKS overexpression effectively uncouples bacterial attachment from pedestal formation, consistent with the observed loss of overlap between Tir and EspF_U_ signals (Fig. S1). These findings are consistent with previous work showing that overexpression of the IRTKS I-BAR or SH3 domains, as well as siRNA knockdown (KD) of IRTKS, decreased EspF_U_-dependent formation of actin rich pedestals (Vingadassalom et al., 2009).

### IRTKS KO reduces Tir-mediated attachment, EspF_U_ recruitment, and pedestal formation

To further investigate the role of IRTKS in Tir membrane localization and Tir-mediated bacterial attachment for pedestal formation, we created an IRTKS knockout (KO) CACO-2BBE cell line using CRISPR/Cas9 genome editing (Fig. S3A). KC12+EspF_U_ infected IRTKS KO cells exhibited significantly lower levels of HA-Tir beneath individual bacterial attachment sites, as quantified using line-scan analysis (Fig. 5A–5C). In many cases, attached bacteria demonstrated little to no HA-Tir enrichment at the point of contact with the apical membrane (Fig. 5B). In a subset of attachments, HA-Tir signal demonstrated strong overlap with the bacterial DAPI signal, suggesting that Tir was retained within surface-bound bacteria (Fig. S4). Bacteria on the surface of IRTKS KO monolayers also demonstrated extensive clustering, as indicated by reduced grid occupancy across confocal image fields (Fig. 5D), and shorter distances between neighboring bacteria (Fig. 5E). These results suggest a reduced capacity for bacterial movement and spreading across the apical surface in the absence of IRTKS, which could be related to defects in KC12+EspF_U_ pedestal formation. To further test this possibility, we examined EspF_U_ recruitment and pedestal formation in IRTKS KO cells. Line scan analysis of pedestals showed a significant decrease of myc-EspF_U_ beneath bacterial attachment sites (Fig. 5F, 5H). The low levels of EspF_U_ still present at these sites indicate that additional pathways likely contribute to EspF_U_ recruitment (Lai et al., 2013; Weiss et al., 2009). Quantification of pedestal formation efficiency revealed a two-fold decrease in IRTKS KO cells (Fig. 5G). In addition, line scan analysis of F-actin intensity revealed a reduction in signal in individual pedestals on the surface of IRTKS KO cells (Fig. 5I). Pedestals that did manage to form on IRTKS KO cells were smaller and demonstrated significantly reduced projected pedestal areas compared to controls (Fig. 5F, 5J). Together these results indicate that loss of IRTKS reduces Tir accumulation beneath bacterial attachment sites, which in turn decreases EspF_U_ recruitment and pedestal assembly, and impairs bacterial spread across the apical surface.

**Figure 5:**
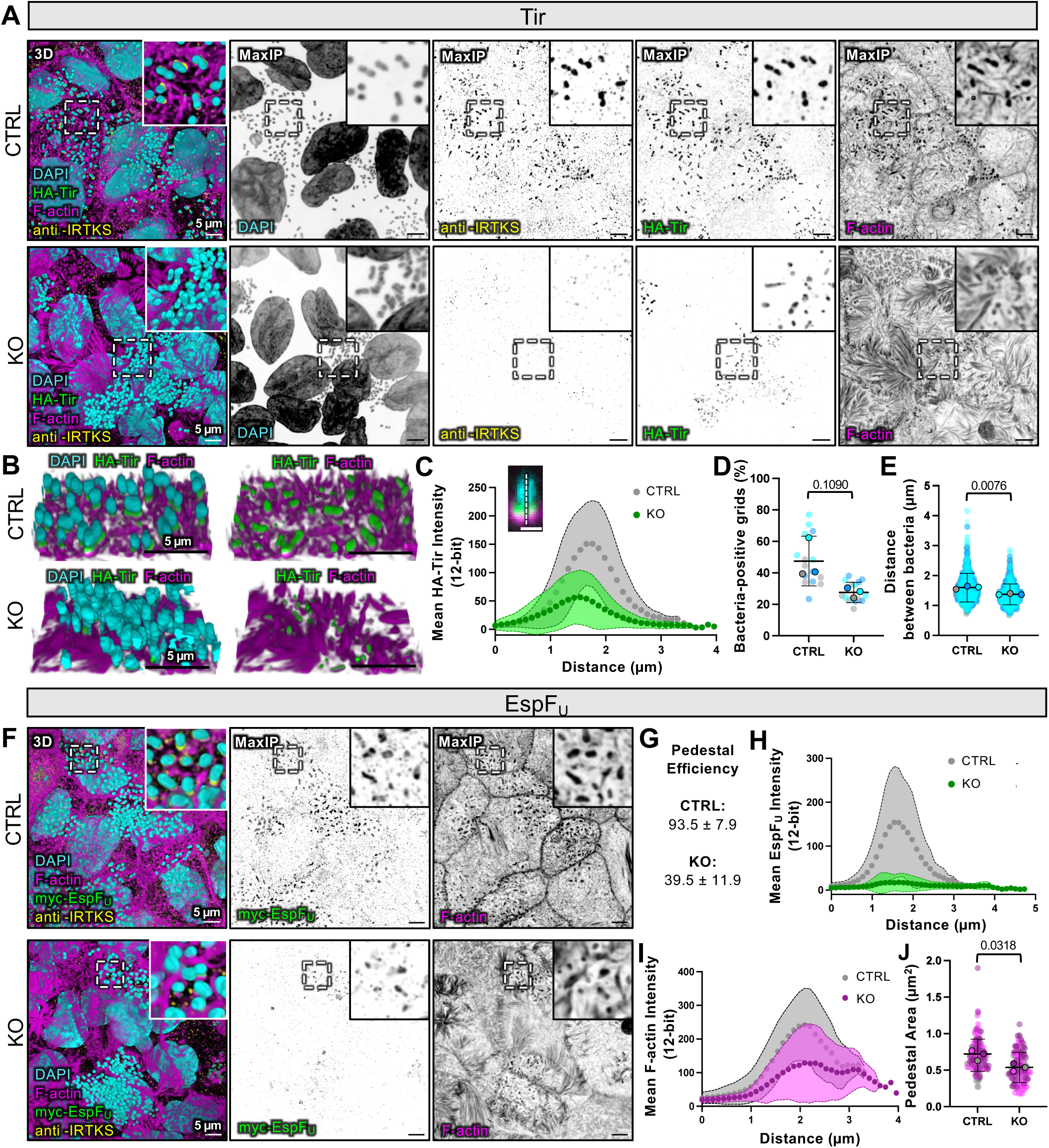
Loss of IRTKS reduces Tir recruitment to the apical membrane and disrupts pedestal formation in CACO-2BBE cells. (A) 3D volume reconstructions and maximum intensity projections (MaxIP) of confocal image volumes of KC12+EspF_U_ infected control and IRTKS KO CACO-2BBE cells stained for DAPI (cyan), HA-Tir (green), IRTKS (yellow), and F-actin (magenta). (B) Tilted view of 3D volume reconstructions of infected control and IRTKS KO CACO-2BBE cells as described in (A), showing the spatial organization of HA-Tir and F-actin at sites of bacterial attachment. (C) Lateral view line scans of individual attached bacteria show the mean HA-Tir fluorescence intensity distribution (12-bit grey values) from control (gray) and IRTKS KO (green) cells; n = 45 pedestals were quantified from three experimental replicates. Mean values are shown as circular points, SD is shown with the transparent color bands. Scale bar represents 1 µm. (D) Percentage of bacteria-positive grids, from fields of view that were subdivided into 5-µm × 5-µm regions; 676 total grids scored from five fields of view per experimental replicate; n = three experimental replicates. (E) Distance between neighboring bacteria (µm), derived from measurements made from 250 pairs of bacteria per experimental replicate; n = three experimental replicates. (F) 3D and MaxIP image volumes of KC12+EspF_U_ infected control and IRTKS KO CACO-2BBE cells stained for DAPI (cyan), myc-EspF_U_ (green), IRTKS (yellow), and F-actin (magenta). (G) Quantification of pedestal formation efficiency, calculated by dividing the number of adherent bacteria with focused F-actin signal by the total counted; 200 adherent bacteria were counted per experimental replicate; n = three experimental replicates. (H-I) Lateral view line scans of individual adherent bacteria showing the mean of (H) myc-EspF_U_ and (I) F-actin fluorescence intensity distribution (12-bit grey values) within control (gray) and IRTKS KO (green) cells; n = 45 pedestals were quantified from three experimental replicates. Mean values are shown as circular points, SD is shown with the transparent color bands. (J) Pedestal area (µm^2^) measurements based on projected F-actin signal area; n = 50 pedestals per condition were quantified per experimental replicate, with n = three experimental replicates. (D, E, J) Data are shown as superplots, where transparent circles represent individual measurements color-coded by experimental replicate, and opaque outlined circles represent the means of each experimental replicate. Statistical comparisons were performed on experimental replicate means. All p-values were calculated with a Welch’s unpaired t-test; exact p-values are shown above brackets. Error bars represent the mean ± SD of the technical replicates. Scale bars represent 5 µm.

To further define the impact of IRTKS loss-of-function on Tir accumulation and pedestal formation, we generated an IRTKS KO HeLa cell line (Fig. 3SB) and examined the impact of KC12+EspF_U_ infection. Characterization of IRTKS KO in this context revealed several features that were similar to the CACO-2BBE IRTKS KO phenotype described above (Fig. 5). While IRTKS KO cells exhibited a similar number of adherent bacteria compared to controls as measured by LPS signal area per cell (Fig. 6A-6C), Tir recruitment beneath bacterial attachment sites and HA-Tir sum intensity were both significantly reduced (Fig. 6B, 6D, 6E). Loss of IRTKS also led to a decrease in myc-EspF_U_ recruitment, reflected in both reduced signal dispersion and sum intensity (Fig. 6H, 6I). Similar to the CACO-2BBE cells, some residual EspF_U_ signal remained, suggesting the presence of additional recruitment pathways. Consistent with these defects, pedestal formation efficiency was significantly reduced in IRTKS KO cells (Fig. 6G), and the pedestals that did form exhibited a smaller area (Fig. 6J). Overall, these results show that IRTKS is required for normal Tir accumulation in the plasma membrane beneath bacterial attachment sites, and for promoting downstream EspF_U_-dependent pedestal formation, independent of host cell surface context/morphology.

**Figure 6:**
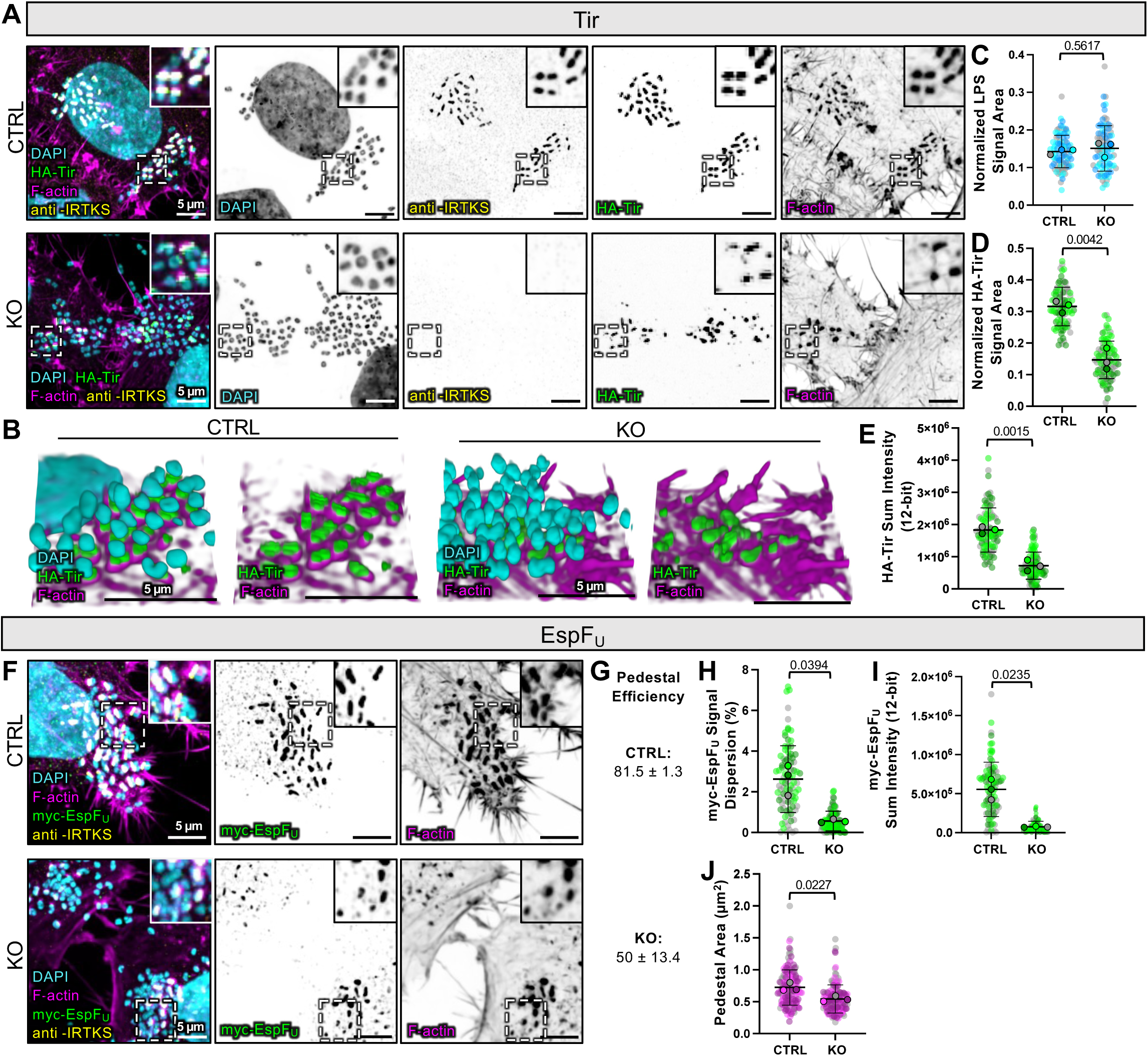
Loss of IRTKS reduces Tir-mediated attachment and pedestal formation in HeLa cells. (A) Maximum intensity projections (MaxIPs) of confocal image volumes of KC12+EspF_U_ infected control and IRTKS KO HeLa cells stained for DAPI (cyan), HA-Tir (green), IRTKS (yellow), and F-actin (magenta). (B) Tilted 3D reconstructions showing infected control and IRTKS KO HeLa cells as described in (A). (C) Quantification of thresholded LPS fluorescence signal area per cell, normalized to total cell area; data were analyzed using LPS-stained confocal images (not shown); 30 cells per condition were quantified per experimental replicate, with n = three experimental replicates. (D) Quantification of thresholded HA-Tir fluorescence signal area, normalized to the LPS signal area; 30 cells per condition were quantified per experimental replicate, with n = three experimental replicates. (E) Sum of HA-Tir 12-bit intensity values per cell from thresholded HA-Tir fluorescence signal; 30 cells per condition were quantified per experimental replicate, with n = three experimental replicates. (F) MaxIP of KC12+EspF_U_ infected control and IRTKS KO cells stained to visualize DAPI (cyan), myc-EspF_U_ (green) IRTKS (yellow), and F-actin (magenta). (G) Quantification of pedestal formation efficiency, calculated by dividing the number of adherent bacteria with focused F-actin signal by the total counted; 200 adherent bacteria were counted per experimental replicate; n = three experimental replicates. (H) Quantification of the percentage of myc-EspF_U_ signal dispersion, from thresholded myc-EspF_U_ fluorescence signal area was normalized to the total cell area; n = 30 cells per condition were quantified per experimental replicate. (I) Sum of 12-bit myc-EspF_U_ intensity values per cell from thresholded myc-EspF_U_ fluorescence signal; 30 cells per condition were quantified per experimental replicate, with n = three experimental replicates. (J) Pedestal area (µm^2^) measurements based on projected F-actin signal area; 50 pedestals per condition were quantified per experimental replicate, with n = three experimental replicates. (C-D, H-J) Data are shown as superplots, where transparent circles represent individual measurements color-coded by experimental replicate, and opaque outlined circles represent the means of each experimental replicate. Statistical comparisons were performed on experimental replicate means. All p-values were calculated with a Welch’s unpaired t-test; exact p-values are shown above brackets. Error bars represent the mean ± SD of the technical replicates. Scale bars represent 5 µm.

### IRTKS is sufficient for promoting Tir membrane insertion and bacterial attachment

Our characterization of IRTKS KO and overexpression phenotypes in CACO-2BBE and HeLa cells collectively suggest that accumulation of Tir in the host plasma membrane depends on IRTKS. However, the current model for pedestal formation positions IRTKS downstream of Tir (Vingadassalom et al., 2009). To clarify the functional relationship between IRTKS and Tir, we sought to determine if IRTKS expression is sufficient to drive the insertion of Tir into the plasma membrane. Here, we assayed Tir membrane insertion by measuring the attachment of *E. coli* strain expressing EHEC intimin alone (*E. coli* + pIntimin^EHEC^) (Campellone et al., 2006); because no other virulence factors are present, pedestals are unable to form and bacterial attachment is a direct readout for the amount of Tir inserted in the membrane. Here, HeLa cells were transfected with plasmids encoding full length EGFP-IRTKS and HA-tagged Tir (pHA-Tir), and then incubated with *E. coli* + pIntimin (Fig. 7A). In the resulting confocal images, we binned cells and scored bacterial attachment as a function of HA-Tir and EGFP-IRTKS intensity levels. Remarkably, cells co-expressing both constructs (pHA-Tir and EGFP-IRTKS) showed a striking increase in bacterial attachment, whereas cells expressing only pHA-Tir or EGFP-IRTKS, or cells that remained untransfected, exhibited little to no bacterial attachment (Fig. 7B, 7C, arrows). Consistent with these observations, a scatter plot of mean IRTKS intensity vs. mean HA-Tir intensity showed that increased expression of both proteins correlated with greater bacterial adherence (Fig. 7D). In contrast to existing pedestal formation pathway models, these findings strongly indicate that Tir insertion into the plasma membrane for intimin binding depends on IRTKS.

**Figure 7:**
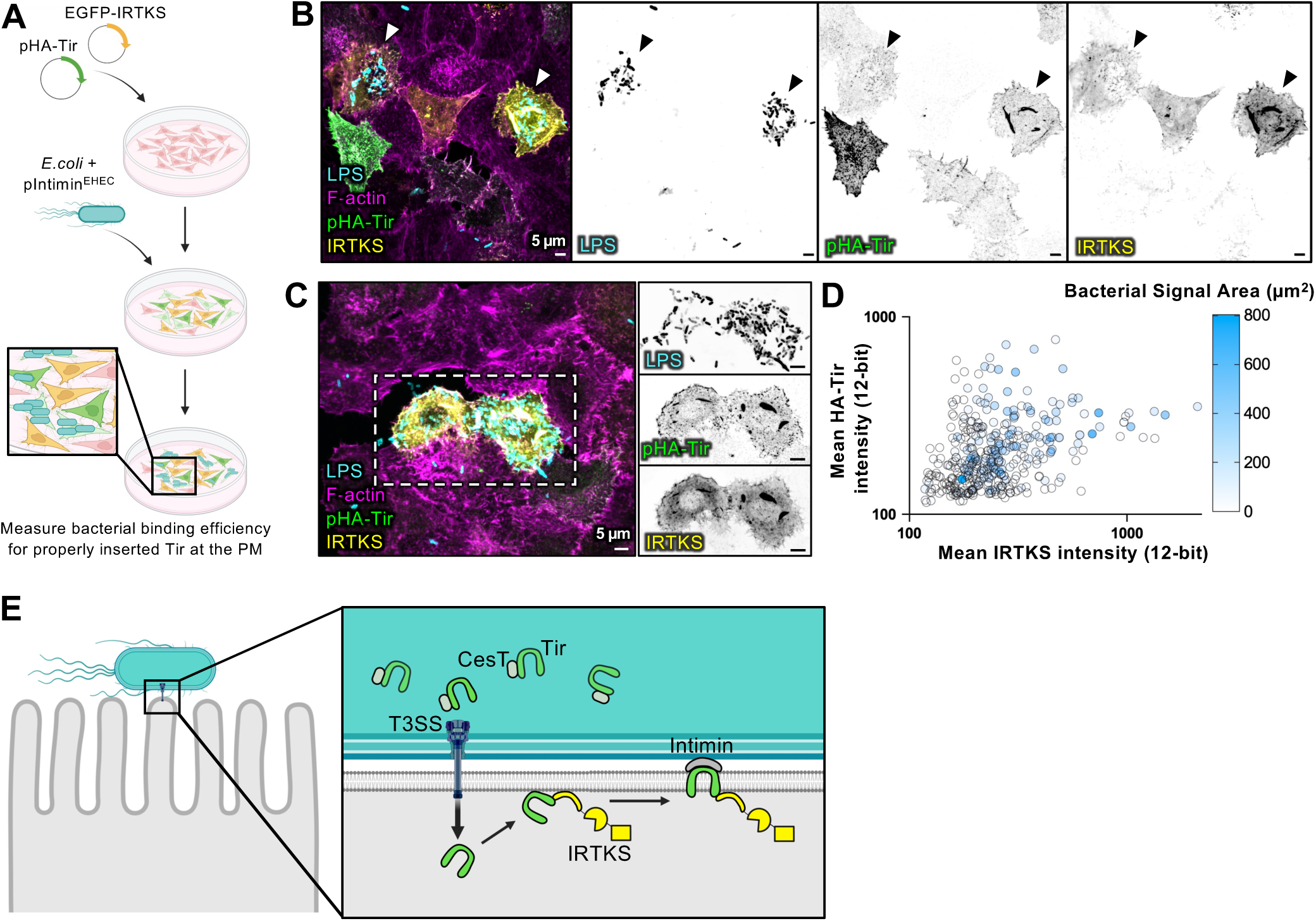
IRTKS promotes Tir membrane insertion for bacterial attachment. (A) Schematic depicting the experimental method used to measure Tir insertion in the plasma membrane. HeLa cells overexpressing pHA-Tir (green) and EGFP-IRTKS (yellow) were incubated with *E.coli*+pIntimin^EHEC^ and bacterial attachment was assayed, as an indicator for proper Tir insertion into the membrane. (B-C) Maximum intensity projections of confocal image volumes of *E.coli*+pIntimin^EHEC^ infection on HeLa cells stained for LPS (cyan), pHA-Tir (green), IRTKS (yellow), and F-actin (magenta). Arrows indicate cells overexpressing both IRTKS and pHA-Tir. (D) Scatter plot of the mean IRTKS intensity (12-bit grey values) versus the mean HA-Tir intensity (12-bit grey values) plotted on a logarithmic scale. Each point represents a single cell, and the color denotes bacterial attachment, measured as bacterial signal area (µm^2^), with darker blue corresponding to an increased attachment; n = 370 cells were quantified from three experimental replicates. All scale bars = 5 µm.

## DISCUSSION

During infection with A/E pathogens, brush border microvilli are remodeled to allow for the formation of actin pedestals on the enterocyte’s apical surface. In the specific case of EHEC, the I-BAR domain containing protein, IRTKS, is hijacked as a scaffolding protein, to link virulence factors Tir and EspF_U_ to form a complex that initiates actin assembly for pedestal formation (Vingadassalom et al., 2009). An important open question in EHEC pathogenesis is how Tir, which is initially injected into the host cell cytosol as a soluble factor, targets to the apical surface and inserts into the plasma membrane to mediate intimate bacterial attachment. In the current study, we sought to test the idea that, by virtue of its Tir- and plasma membrane-binding abilities, host factor IRTKS facilitates Tir membrane insertion upstream of its role as a scaffold for EspF_U_. In our experiments, we leveraged multiple new gain- and loss-of-function models and used high performance microscopy to assay the localization and organization of pedestal components. Collectively, our findings confirm that IRTKS is necessary for normal EspF_U_-dependent pedestal formation (Vingadassalom et al., 2009), and further suggest that this factor functions early in the pedestal formation pathway, to promote Tir accumulation in the host cell plasma membrane.

We initially predicted that overexpression of IRTKS would enhance pedestal formation by enabling Tir to indirectly recruit more EspF_U_ to the apical surface. In contrast, we unexpectedly observed defects in pedestal formation, characterized by extensive Tir dispersion across the plasma membrane. Additionally, cells overexpressing IRTKS also exhibited higher levels of total Tir. This was paralleled by a striking increase in bacterial adherence, suggesting that dispersed Tir was functionally inserted into the plasma membrane and capable of supporting intimate attachment. We further found that the I-BAR domain, which binds to Tir and the plasma membrane, is required for Tir dispersion; a construct lacking an intact I-BAR (ΔI-BAR) was unable to drive this phenotype, whereas constructs containing this motif (I-BAR only and SH3*) exhibited dispersion levels that trended strongly upward.

How might IRTKS overexpression exert these effects on Tir? Based largely on studies of EPEC Tir, current models posit that, after injection through the T3SS along with chaperone CesT, Tir is inserted into the host cell plasma membrane as a hairpin loop (Allen-Vercoe et al., 2005; Luo et al., 2000; Thomas et al., 2005). In this conformation, both N- and C-termini extend into the host cytoplasm, whereas the central loop extends into extracellular space to participate in intimin binding (Fig. 1A) (Kenny, 1999; Kenny et al., 1997). Although the mechanistic details of how Tir accumulates in the plasma membrane following its injection into host cells have remained unclear, previous work showed that ectopically expressed Tir can insert into the plasma membrane and bind intimin (Campellone et al., 2006), and *in vitro* studies found that purified Tir can insert directly into lipid vesicles (Race et al., 2006). Both of those findings suggest that additional bacterial effectors are not required for Tir membrane insertion. However, when we ectopically co-expressed Tir with IRTKS in HeLa cells, and probed membrane insertion using an *E. coli* strain expressing EHEC intimin (*E. coli* + pIntimin^EHEC^), cells containing both IRTKS and Tir demonstrated much higher levels of bacterial adherence relative to cells expressing IRTKS or Tir alone. These results indicate that IRTKS directly promotes the functional insertion of Tir into the plasma membrane, above basal levels.

IRTKS is well equipped for driving Tir membrane insertion early in the pedestal formation pathway, as it contains an N-terminal I-BAR domain that binds to both the C-terminal cytoplasmic tail of Tir and the plasma membrane (Millard et al., 2007; Postema et al., 2018; Vingadassalom et al., 2009). Whether the I-BAR can bind Tir and membrane simultaneously remains unknown, although the amino acids (L27, K107, R192, F195) that are predicted to interact with the C-terminal NPY motif of Tir (de Groot et al., 2011) do not reside in the basic patches that are expected to support lipid binding (Mattila et al., 2007; Millard et al., 2005). Based on our new findings and these points, we propose an updated working model whereby Tir is injected into the host cell cytoplasm, and through its C-terminal tail it binds directly to the plasma membrane-associated I-BAR domain of IRTKS (Fig. 7E). Anchoring Tir at the inner leaflet in this way could potentially accelerate its insertion into the plasma membrane and in turn promote intimate bacterial adherence.

Overexpression of IRTKS also significantly disrupted pedestal formation, which provided clear evidence that - at high IRTKS concentrations - bacterial attachment can be uncoupled from downstream F-actin assembly. Staining for EspF_U_ revealed that this factor was dispersed, with a lack of focused enrichment beneath adherent bacteria, providing a mechanistic explanation for pedestal failure. Uncoupling between bacterial attachment and pedestal formation suggests that the normal, functional stoichiometry of the Tir-IRTKS-EspF_U_ complex (1:1:1) was disrupted in IRTKS overexpressing cells. Because IRTKS is the central, bivalent link that simultaneously binds to Tir and EspF_U_, the formation of 1:1:1 tripartite complexes will be compromised at high concentrations of IRTKS. Under these conditions, Tir and EspF_U_ would become physically uncoupled due to the formation 1:1 binary complexes of Tir-IRTKS and IRTKS-EspF_U_, leading to the phenotypes we observed here. Our findings are also consistent with previous work showing that overexpression of IRTKS truncation constructs disrupted pedestal F-actin assembly (Vingadassalom et al., 2009).

Loss of IRTKS in both CACO-2BBE and HeLa cells reduced Tir accumulation at the bacteria-host interface. Although overall bacterial adherence was not reduced, IRTKS KO reduced Tir signal relative to adherent bacteria, suggesting impaired Tir-mediated attachment and indicating that IRTKS is required for normal Tir accumulation in the plasma membrane. IRTKS KO also reduced F-actin enrichment in pedestals, which led to impaired bacterial spread, as reflected by increased bacterial clustering. These findings are consistent with previous work showing that pedestal F-actin turnover drives bacterial surface motility and in turn promotes cell-to-cell spread (Velle and Campellone, 2017). These results are also aligned with earlier studies showing that siRNA-mediated knockdown of IRTKS decreases EspF_U_-dependent pedestal formation (Vingadassalom et al., 2009). Thus, IRTKS is required for normal Tir accumulation in the plasma membrane early in the pedestal formation pathway, as well as the cytoskeletal remodeling needed for efficient pedestal formation and bacterial spread.

Under normal conditions, IRTKS is found at the distal tips of microvilli and is implicated in microvillar elongation (Gaeta et al., 2021; Postema et al., 2018). During infection, this factor is repurposed by EHEC to promote pedestal formation (Vingadassalom et al., 2009). In this work, we found that, beyond its function as a scaffolding link between Tir and EspF_U_, IRTKS plays an important upstream role in promoting Tir accumulation in the plasma membrane, to support intimate bacterial attachment. Several open questions remain; perhaps most importantly, the biochemical details of how IRTKS enhances the functional insertion of Tir into the plasma membrane remain to be determined. *In vitro* assays that employ synthetic vesicles for well-defined targets for Tir insertion (Race et al., 2006) could be leveraged to delineate biochemical mechanisms underpinning the impact of IRTKS. Additionally, while IRTKS KO cells exhibited defects in pedestal formation, pedestals are still able to form in these models, which is consistent with a previous study (Vingadassalom et al., 2009), indicating that other undefined pathways may contribute to pedestal assembly. The IRTKS KO cell lines generated and examined in this study would provide powerful models for defining these alternative mechanisms and extending our understanding of EHEC pathogenesis.

## MATERIALS AND METHODS

### Bacterial strains

Bacterial strains used in these studies were (i) KC12+EspF_U_ and (ii) *E. coli* + pIntimin (generous gifts from the Campellone Lab) (Campellone et al., 2006; Campellone et al., 2002). Prior to infections, *E. coli* strains were cultured in Luria-Bertani (LB) broth and grown in a shaking incubator at 37° C overnight in the presence of selective antibiotics (30 mg/ml Kan and 50 mg/ml Amp for KC12+EspF_U_; 50 mg/ml Amp for *E. coli* + pIntimin). The next day, cultures were diluted 1:25 in Gibco Dulbecco’s Modified Eagle Medium: Nutrient Mixture F12 (DMEM/F12) and these starter cultures were grown at 37°C with 5% CO_2_ for 3 hrs, to enhance the expression of T3SS effectors.

### Mammalian cell culture

HeLa cells (generous gift from Dr. Ryoma Ohi) and HEK293T cells (purchased from ATCC #CRL-3216) were maintained as subconfluent cultures in Dulbecco’s Modified Eagle’s Medium (DMEM) with high glucose, 2 mM L-Glutamine (Corning #10-013-CV), 1% L-glutamine (Corning #25-005-CI), and 10% fetal bovine serum (FBS) (R&D Systems). CACO-2BBE cells (ATCC #CRL-2102) were maintained in subconfluent monolayers using the same medium supplemented with 20% FBS. To create polarized CACO-2BBE monolayers for infection, cells were grown to confluency on glass coverslips in 6-well plates, and their media was changed every 2-3 days for at least 21 days post confluency. All cells were maintained at 37° C with 5% CO_2_ and regular mycoplasma testing was performed using the MycoStrip Mycoplasma Detection Kit (InvivoGen #rep-mys-50).

### Infection assays

HeLa cells were plated 24 hrs before infection at 5×10^5^ cells/well to obtain 70-90% confluency on glass coverslips in 6-well plates. On the day of infection, cells were washed with PBS and the media was switched to DMEM/F12. Cells were then infected with a dilution of the starter cultures described above; CFU counts were performed to obtain a range of 0.35-0.40. For HeLa cell KC12+EspF_U_ infections, cells were infected for 3 hrs; 1 hr post infection, cells were washed once with PBS to remove unbound bacteria, fresh DMEM/F12 media was added, and cells were incubated for an additional 2 hrs. For *E. coli* + pIntimin binding assays, cells were infected for 5 hrs, with fresh media added 3.5 hrs post infection. Before infecting polarized CACO-2BBE cells, cells were washed once with PBS, and media was replaced with DMEM/F12. CACO-2BBE cells were infected with KC12+EspF_U_ for 6 hrs. 1 hr post infection, cells were washed with PBS to remove unbound bacteria, fresh DMEM/F12 media was added, and cells were allowed to incubate for an additional 5 hrs. All infections were performed at 37° C and 5% CO2.

### Plasmid transfections

HeLa cells were transfected 24 hrs before infection at 8×10^5^ cells/well in 6-well plates. Transfections were performed using Lipofectamine 2000 (Thermo Fischer #11668019) according to the manufacturer’s protocol. Cells were incubated in Lipofectamine for 4-6 hrs and then left overnight in fresh DMEM. IRTKS overexpression experiments were performed with the following plasmids: pEGFP-C1-IRTKS (a.a. 1-511), I-BAR alone (a.a. 1-249), ΔI-BAR (a.a. 250-511), IRTKS-SH3* construct with mutations W378K and W391K. These constructs were created and validated by Dr. Meagan Postema as previously described (Postema et al., 2018). For ectopic expression of Tir, a pHA-Tir plasmid (generous gift from Dr. Kenneth Campellone) was used as previously described (Campellone et al., 2006). The identity of all plasmids used in this study was validated by sequencing.

### IRTKS knock-out (KO) cell line generation

CACO-2BBE and HeLa IRTKS KO cell lines were generated with the Lenti-CRISPR v2 system (Addgene #52961), using gRNAs designed to target exon 3 of human IRTKS genomic sequence [Exon 3 Fwd CACCGAGTTGACACGGGGGACCCAG; Exon 3 Rev AAACCTGGGTCCCCCGTGTCAACTC]. PCR and sequencing primers targeted regions surrounding Exon 3, where Cas9 was predicted to generate a double-strand break [Exon 3 Fwd GACTTTCAGCACCTGCTGCT; Exon 3 Rev AGTCACAGGAAACACTCTGGCCA]. To generate the IRTKS KO in HeLa cells, WT HeLa cells were transfected with Lipofectamine containing the Lenti-CRISPR v2 gRNA plasmid with the gRNAs targeting IRTKS exon 3. 48 hrs post-transfection, cells were placed under puromycin selection (1 mg/ml) for 6 days. To obtain single clones, puromycin-resistant cells were isolated with fluorescence-activated cell sorting, and clonal isolates were expanded in 96-well plates under standard growth conditions.

To generate IRTKS KO in CACO-2BBE cells, parent line cells were transduced with lentivirus that contained the Lenti-CRISPR v2 gRNA plasmid. Lentivirus was generated by transfecting HEK293T cells (5×10^5^ cells/well) with the Lenti-CRISPR v2/gRNA plasmid, pMD2.G envelope plasmid (Addgene #12259), and psPAX2 packaging plasmid (Addgene #12260) using PEI (Polyscience #23966-100). For efficient lentiviral production, cells were incubated for 48 hrs before collecting the lentivirus-containing media. Parent line CACO-2BBE cells were then transduced with lentivirus particles supplemented with polybrene (Sigma #TR-1003-G). 48 hrs post-transduction, cells were placed under puromycin selection (10 mg/ml) for 2 weeks. Serial dilutions were performed to obtain single clone populations. Single clones were validated by western blotting.

### Western blotting

To create lysates for western blot analysis, cells were washed with cold PBS, scraped, and incubated on ice in Cell Lytic lysis buffer (Sigma #C2978) containing protease inhibitors (Complete, Roche # 05892791001), Pefabloc, and ATP. Cell lysates were mixed with SDS-PAGE sample buffer containing 2-mercaptoethanol (Sigma #M3148) and boiled for 5 min. Samples were then separated by SDS-PAGE (NuPAGE TM Bis-Tris Gel NP0323BOX, Invitrogen) and gels were then transferred to nitrocellulose membranes using a TransBlot Turbo Transfer System (BIO-RAD). Membranes were blocked with 5% BSA and incubated with primary antibodies in 3% BSA overnight at 4°C. The next morning, membranes were washed 3 times with PBS-T (1X PBS and 0.1% Tween-20) and then incubated with secondary antibodies for 1 hr at RT in the dark. Proteins were detected by fluorescence using a BIO-RAD ChemiDoc system. Antibodies used in this study are listed in the Key Resource Table.

### Immunofluorescent staining

After infection, all cells were washed once with PBS and fixed with 4% paraformaldehyde (Electron Microscopy Sciences #15710) for 15 min at 37° C. Cells were then rinsed 3 times, 5 min each, in PBS and permeabilized with 0.1% Triton X-100 (Sigma # T8787) in PBS for 15 mins at RT. After permeabilization, cells were rinsed 3 times, 5 minutes each in PBS, and blocked with 5% BSA (Research Products International #9048-46-8) in PBS for 1 hr at 37° C. Cells were then rinsed once with PBS and the primary antibody (diluted in 1% BSA) was added for 1 hr at 37°C. After primary antibody incubations, cells were washed 4 times, 5 min each, with PBS. Secondary antibody and phalloidin for labeling F-actin (diluted in 1% BSA) was then added for 1 hr at RT in the dark. DAPI (1:100; Millipore Sigma #D9542) was added after the secondary antibody for 15 min at RT in the dark. Finally, cells were washed 4 times, 5 minutes each with PBS and coverslips were mounted on glass slides with ProLong Gold (Invitrogen #P36930). Antibodies used in this study are listed in the Key Resource Table.

### Light microscopy

Stained samples were imaged using a spinning disk confocal microscope consisting of a Nikon Ti2 inverted light microscope with a Yokogawa CSU-W1 spinning disk confocal head, a Photometrics Prime 95B sCMOS camera, four excitation LASERs (405, 488, 568, and 647 nm), and a 100X/1.49 NA Apo TIRF oil immersion objective. Imaging acquisition parameters were matched across samples during image acquisition.

### lmage analysis

Unprocessed images were used to analyze data. To improve visualization for figure preparation, maximum intensity projections (MaxIPs) and 3D reconstructed confocal images were denoised and deconvolved using Nikon Elements software. Image channels were merged, separated, and cropped using FIJI. Preceding analysis of fluorescence intensities, raw images were manually thresholded using the “Binary” menu “Define Threshold” function in Nikon Elements, without applying any additional filtering. Thresholds were defined based on the relevant fluorescence signal - LPS, HA-Tir, phalloidin/F-actin, or myc-EspFU - to minimize background fluorescence. Quantification was then performed on the thresholded regions encompassing the signal of interest, as described below.

#### Line scans for IRTKS and other key pedestal components

For HeLa cells, lines scans measuring mean fluorescence intensity of DAPI, HA-Tir, IRTKS, myc-EspF_U_, ArpC2, and phalloidin (F-actin) signals in pedestals were generated using MaxIPs of confocal volumes, by drawing lines from the distal end of the DAPI signal to the basal end of the F-actin signal using Fiji. For Caco-2BBE cells, line scans were drawn on lateral views. For each plot, distance and fluorescence intensity were normalized from 0 to 1, where 0 represents the minimum value in the dataset and 1 represents the maximum value. Each fluorescence channel was normalized independently, and normalized profiles were then plotted together.

#### HA-Tir and myc-EspF_U_ Signal Dispersion and Sum Intensity

MaxIPs of confocal volumes of control cells or cells overexpressing IRTKS or mutant constructs, were thresholded to define the area occupied by HA-Tir and myc-EspF_U_ fluorescence signals; this value was then normalized to the total cell area. Thresholded areas were also used to measure the sum intensity of HA-Tir and myc-EspF_U_ signals.

#### LPS Signal Volume and Area

To measure LPS Signal volume, 3D confocal volumes of control cells or cells overexpressing IRTKS or mutant constructs were subject to thresholding, to define the LPS signal volume. LPS signal volumes were then normalized to total cell area. To quantify LPS Signal Area, MaxIPs of confocal volumes of control or IRTKS KO cells were thresholded to define the area occupied by LPS signal, and this value was then normalized to total cell area.

#### HA-Tir Signal Area

MaxIPs of confocal volumes of control or IRTKS KO HeLa cells were subject to thresholding to capture the area occupied by HA-Tir fluorescence signal. HA-Tir signal area was then normalized to LPS signal area measured from the same cell.

#### Percentage of Bacteria-positive Grids

MaxIPs of confocal volumes were subdivided into 5-µm × 5-µm grids, with a total of 676 grids per field. Grids containing bacteria-positive signal were manually counted and expressed as a percentage of the total number of grids per field.

#### Number of Pedestals per Cell and Pedestal Formation Efficiency

MaxIPs of confocal volumes of control cells and cells overexpressing IRTKS or mutant constructs were used to manually count the number of pedestals per cell. Pedestal formation efficiency for control and IRTKS KO cells was determined by dividing the number of F-actin-dense pedestals by the total number of adherent bacteria. For each experimental replicate, 200 adherent bacteria were scored.

#### Pedestal Area

For control and IRTKS KO cells, regions of interest were manually drawn around the F-actin signal associated with a pedestal-forming bacterium; this region was then used to measure pedestal signal area.

#### Pearson Correlation Coefficient Analysis

MaxIPs of confocal volumes of control cells or cells overexpressing IRTKS mutant constructs were subject to thresholding, to define myc-EspF_U_ and TOMM20 fluorescence signal areas. Pearson correlation coefficient analysis between myc-EspF_U_ and TOMM20 signals was performed using Nikon Elements.

### Statistical analysis

All graphs were generated and statistical analyses performed using GraphPad Prism v10.0. Statistical comparisons were performed using experimental replicate means (i.e. the larger points on Superplots). For comparisons between two groups, p-values were calculated using Welch’s unpaired t-test. For comparisons among multiple groups, p-values were calculated using ordinary one-way ANOVA with multiple comparisons.

**TABLE 1.**
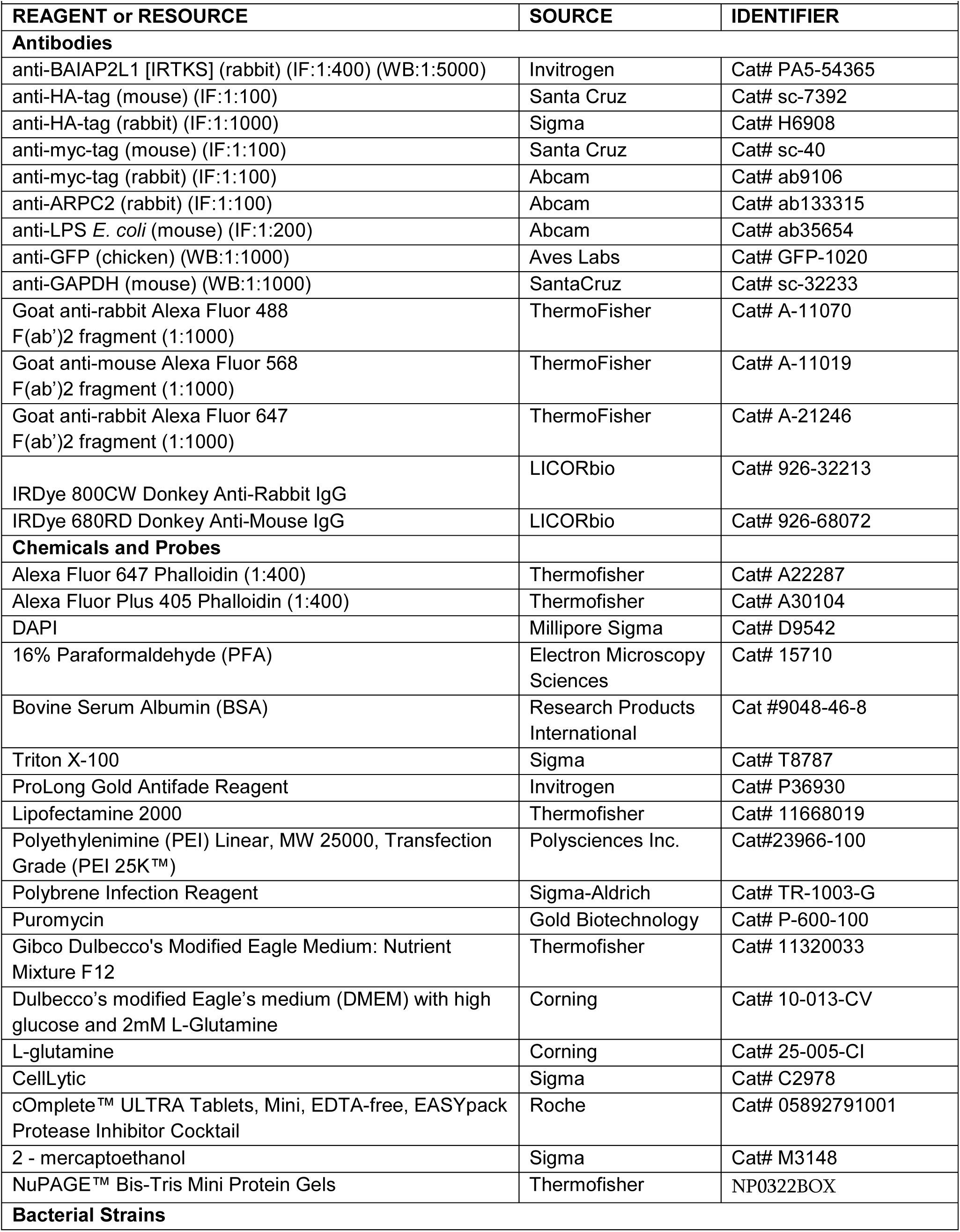
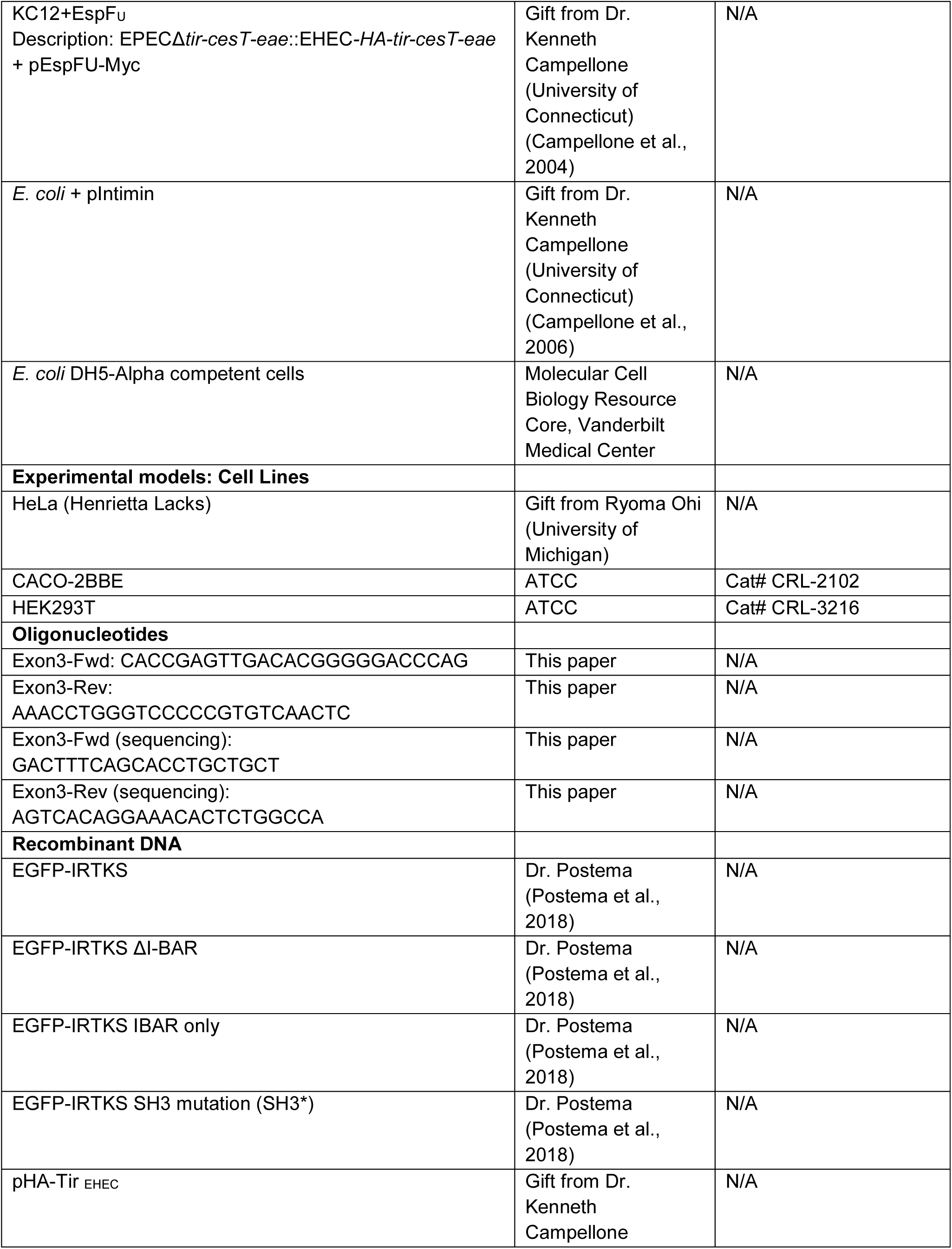
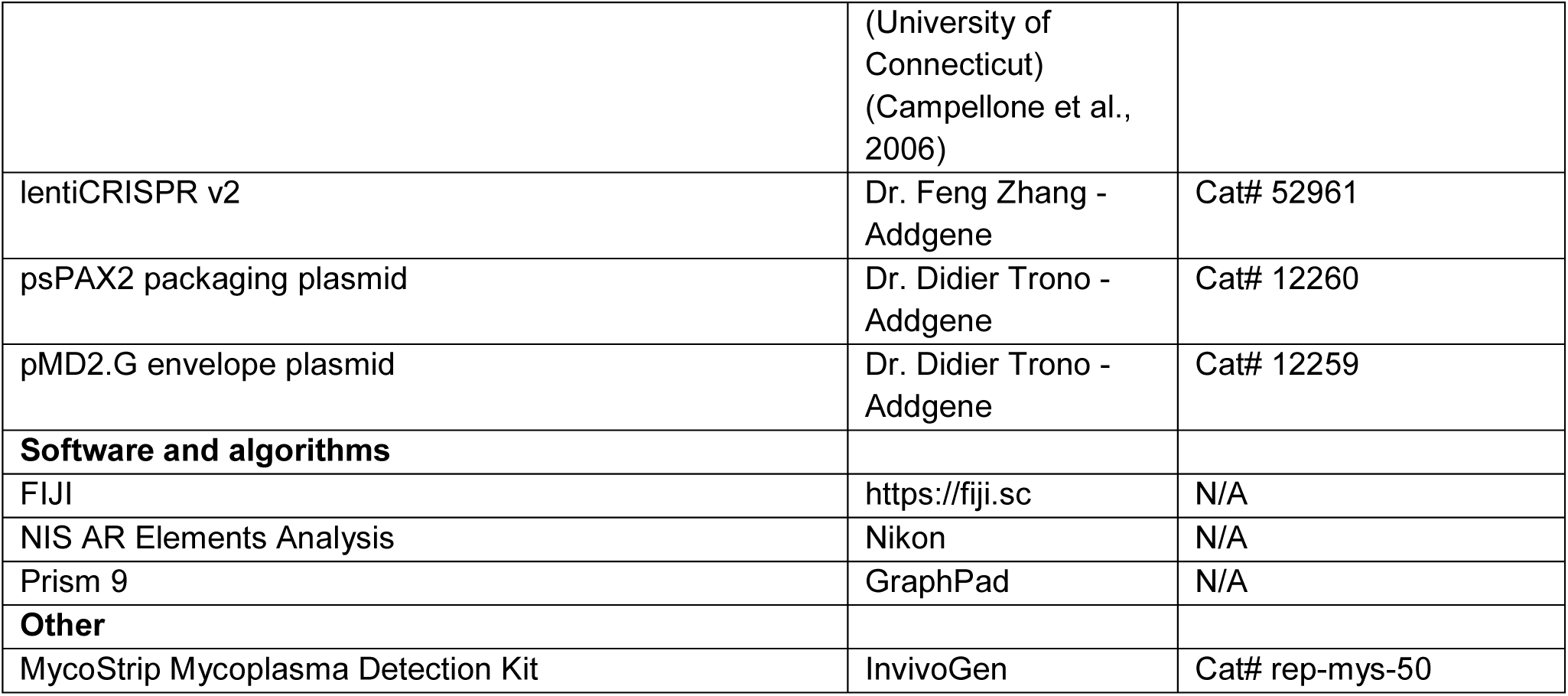
KEY RESOURCES.

## ACKNOWLEDGEMENTS

The authors would like to thank all members of the M.J.T. laboratory for their feedback and guidance. We thank Dr. Kenneth Campellone for supplying plasmids and bacterial strains used in this study. This work was supported by NIH grants R01-DK095811 and R01- DK111949 (M.J.T.). We also acknowledge the Lacks family and are grateful for the use of HeLa cells, which heavily contributed to the discoveries in this work.

## AUTHOR CONTRIBUTIONS

Conceptualization, J.B. and M.J.T.; methodology, J.B. and M.J.T.; validation, J.B.; formal analysis, J.B., F.Z.M., and M.J.T.; investigation, J.B., F.Z.M., and C.I.R.; writing, J.B. and M.J.T.; visualization, J.B.; supervision, M.J.T.; project administration, M.J.T.; funding acquisition, M.J.T.

## Abbreviations

EHEC: Enterohemorrhagic *Escherichia Coli*
EPEC: Enteropathogenic *Escherichia Coli*
A/E: Attaching and Effacing
LEE: Locus of Enterocyte Effacement
T3SS: Type III Secretion System
Tir: Translocated Intimin Receptor
EspF_U_: *Escherichia coli* Secreted Protein F in Prophage U
IRTKS: Insulin Receptor Tyrosine Kinase Substrate
I-BAR: Inverse-Bin-Amphiphysin-Rvs
IRSp53: Insulin Receptor Tyrosine Kinase Substrate p53
SH3: SRC Homology 3
DPC: days post-confluency
LPS: Lipopolysaccharide

## Notes

### Competing Interest Statement

The authors have declared no competing interest.

