## Supplemental Figure for "IRTKS promotes Tir membrane insertion for intimate bacterial attachment and subsequent pedestal formation"

By Burgos-Rivera et al.

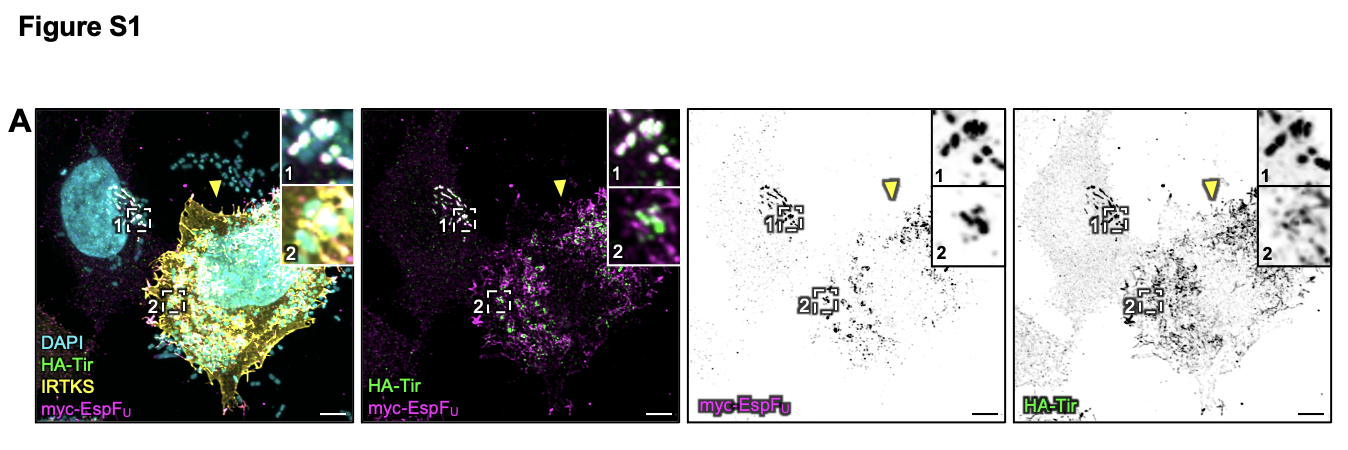

**Figure S1: High overexpression of IRTKS leads to uncoupling of Tir and EspF_U_**. (A) Maximum intensity projection of confocal image volume of KC12+EspF_U_ infected HeLa cells overexpressing IRTKS. Stained for DAPI (cyan), HA-Tir (green), IRTKS (yellow), and myc-EspF_U_ (magenta). Zoom 1 shows a cell expressing low levels of IRTKS and Zoom 2 shows a cell expressing higher levels (yellow arrow). Scale bar = 5 µm.

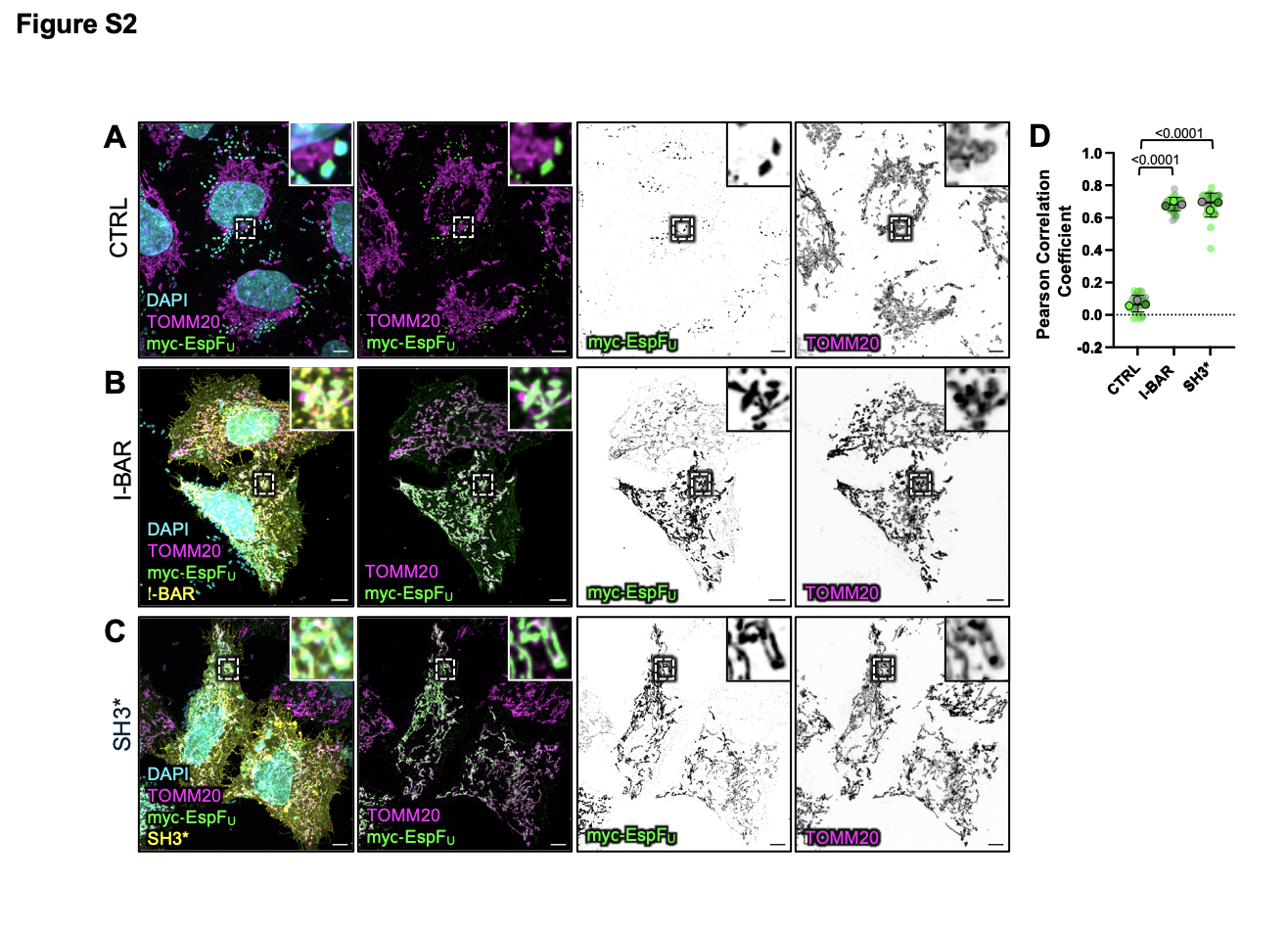

**Figure S2: Deletion or mutation of the IRTKS SH3 domain mislocalizes EspF_U_ to the mitochondria.** (A-C) Maximum intensity projections of confocal image volumes of KC12+EspF_U_ infected (A) control HeLa cells and cells overexpressing (B) EGFP-IRTKS I-BAR, or (C) EGFP-IRTKS SH3*; signals are DAPI (cyan), myc-EspF_U_ (green), and TOMM20 (magenta). Scale bars = 5 µm. (D) Quantification of the Pearson Correlation Coefficient of thresholded myc-EspF_U_ and TOMM20 fluorescence signals. Data is shown as a superplot with n = 3 experimental replicates. Transparent circles represent 10 cells per condition, color-coded by experimental replicate, while opaque outlined circles represent the mean of each experimental replicate. Statistical comparisons were performed on experimental replicate means. All p-values were calculated with an ordinary one-way ANOVA with multiple comparisons. Error bars represent the mean ± SD of the technical replicates.

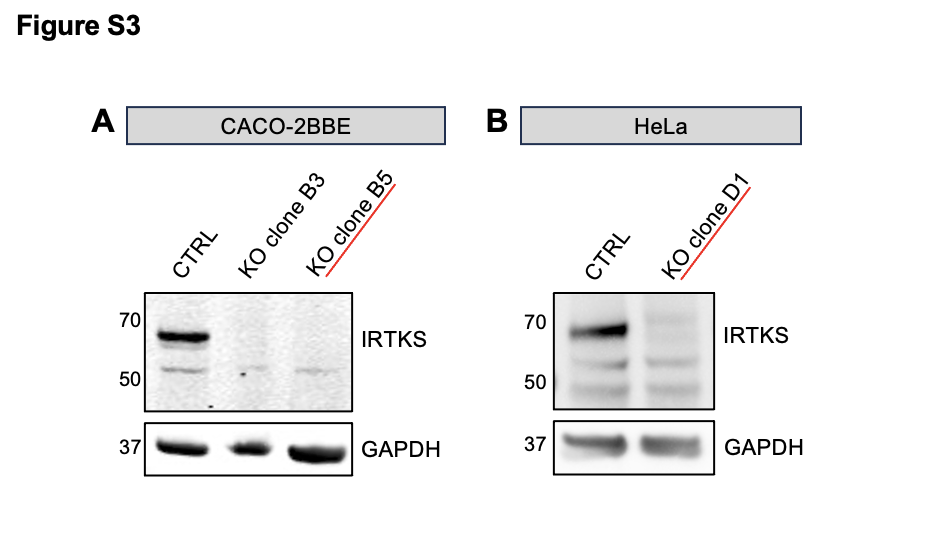

**Figure S3: Validation of IRTKS KO in genome-edited HeLa and CACO-2BBE cell line**s. (A) Cell lysates from control and IRTKS KO CACO-2BBE single clones B3 and B5 were immunoblotted for IRTKS. The B5 clone was selected for experiments shown in this study. (B) Cell lysates from control and IRTKS KO HeLa cells single clone D1 were immunoblotted for IRTKS. GAPDH was used as a loading control in both cases.

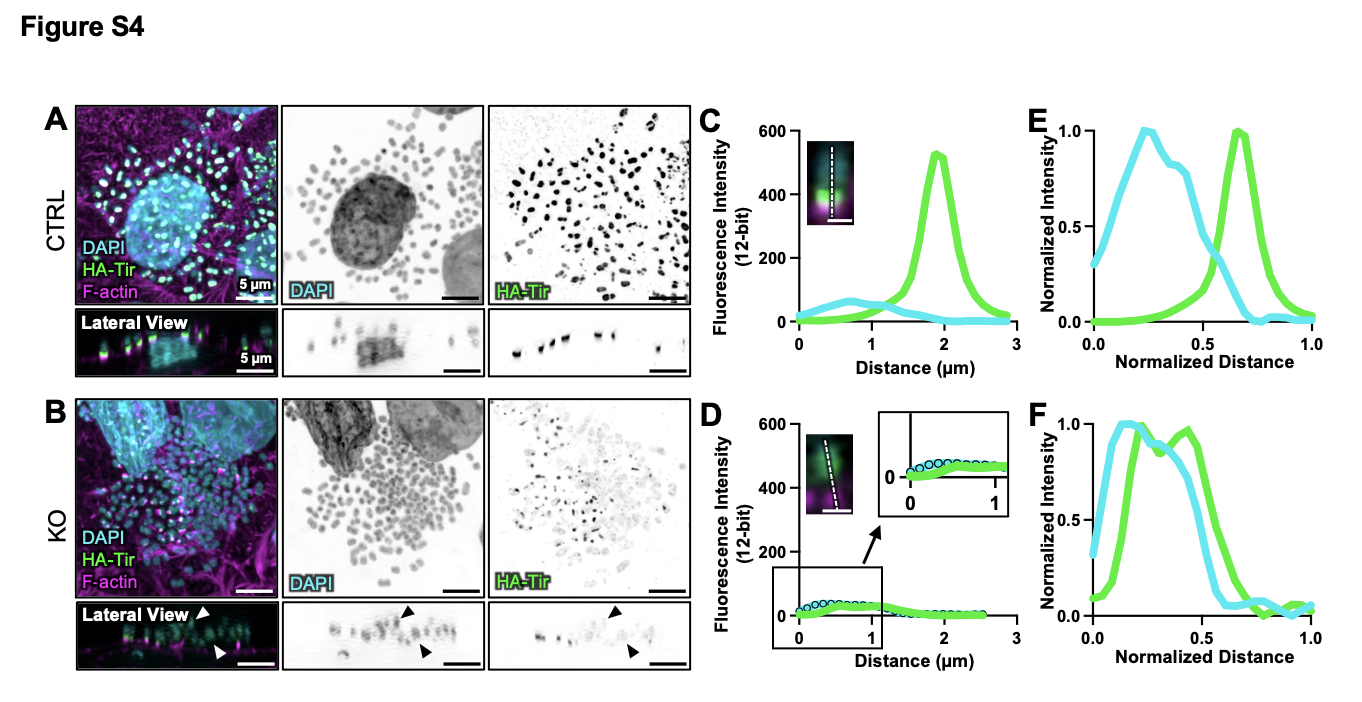

**Figure S4: Tir signal is retained inside bacteria.** (A-B) Maximum intensity projections of confocal image volumes of KC12+EspF_U_ infected control and IRTKS KO CACO-2BBE cells stained for DAPI (cyan), HA-Tir (green), and F-actin (magenta). Lateral views show one z-slice. Scale bars = 5 µm. (C-D) Line scan plots showing the fluorescence intensity (background-subtracted) distribution and localization of DAPI (cyan) and HA-Tir (green) of a single adherent bacterium in (C) control and (D) IRTKS KO CACO-2BBE cells. The lateral view image shows a representative line drawn to generate the plot. Scale bars = 1 µm. Zoom inset in (D) shows overlapping signals. (E-F) Normalized line scan plots corresponding to (C) and (D), respectively.
